# Differential kinetic analysis using nucleotide recoding RNA-seq and bakR

**DOI:** 10.1101/2022.09.02.505697

**Authors:** Isaac W. Vock, Matthew D. Simon

**Author notes:** Corresponding author Author emails: Isaac Vock, Matthew Simon.

## Abstract

Conventional RNA sequencing (RNA-seq) provides limited information about the kinetic mechanisms underlying changes in RNA levels. Nucleotide recoding RNA-seq methods (NR-seq; e.g., TimeLapse-seq, SLAM-seq, etc.) are widely used approaches to identify changes in RNA synthesis and degradation kinetics, yet no software exists to rigorously compare the parameters of RNA kinetics between experimental conditions. We developed bakR to address this need. bakR relies on Bayesian hierarchical modeling of NR-seq data to increase statistical power by sharing information across transcripts. Using simulated and real data, we validate bakR and demonstrate how it provides new insights into the kinetics of RNA metabolism.

## Background

Regulation of gene expression at the RNA level is achieved by regulating rates of RNA synthesis and degradation (1, 2). An increase in the abundance of an RNA transcript could result from an increase in the transcript’s synthesis rate or a decrease in its degradation rate (i.e., transcript stabilization). Therefore, understanding the mechanisms that underly gene expression regulation requires methods to identify changes in the kinetics of RNA metabolism as well as changes in RNA levels.

While standard RNA-seq can identify changes in RNA levels, specialized sequencing-based techniques are required to rigorously investigate the kinetic causes of those changes (3). Metabolic labeling with nucleotide analogs followed by enrichment and sequencing of labeled RNA is one such strategy (4–8). Further innovation was needed because these enrichment-based techniques require substantial amounts of starting RNA, introduce biochemical biases during enrichment, and cannot distinguish the desired enriched RNAs from non-trivial levels of contamination (9, 10). Previously, our lab and others addressed this shortcoming by developing nucleotide recoding RNA-seq methods (NR-seq; Fig. 1A). These techniques combine s^4^U metabolic labeling and nucleotide-recoding chemistry (TimeLapse (11), SLAM (12), TUC (13), etc.) to either convert or disrupt the hydrogen bonding pattern of incorporated s^4^U. This yields apparent T-to-C mutations in the RNA-seq data that indicate sites of s^4^U incorporation and can be used to estimate the fraction of extracted RNA that was synthesized after the introduction of metabolic label (hereafter, fraction new) (14, 15). This adds kinetic information to the snapshot provided by RNA-seq while eliminating the need for enrichment of labeled RNA.

**Fig. 1:**
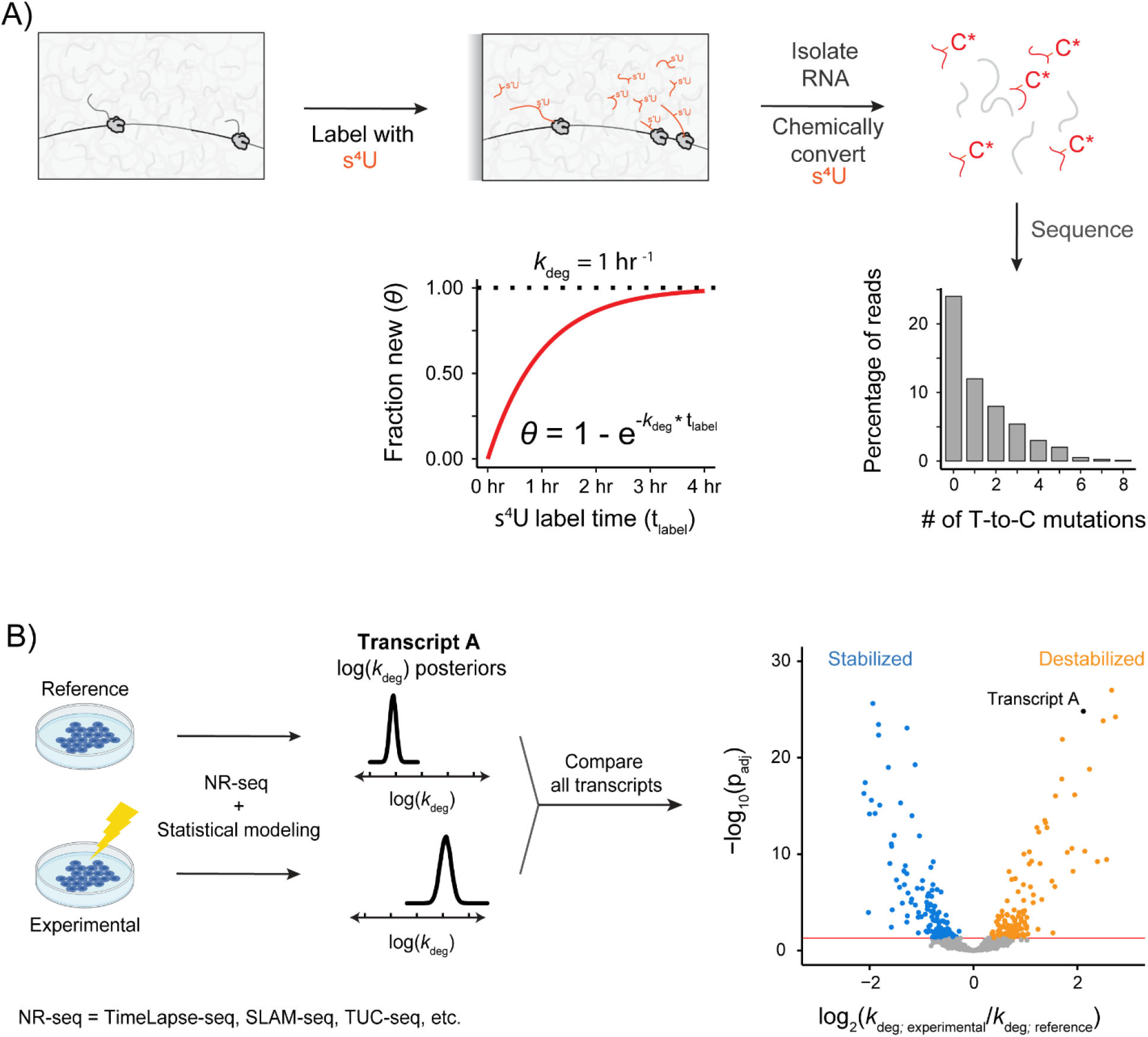
NR-seq can be used to identify differential RNA kinetics. **A)** Schematic of a typical NR-seq experiment. At steady-state, the fraction new has a simple relationship with the degradation rate constant. **B)** Statistical modeling provides estimates of RNA kinetic parameters. In this anecdotal example, Transcript A is less stable in the Experimental condition than in the Reference condition.

NR-seq provides an approach to perform differential kinetic analyses to complement differential expression analyses (Fig. 1B). Rigorous differential kinetic analysis requires a statistical tool that compares replicates of NR-seq data collected in two or more different biological conditions. Such a tool would have to be robust to the low number of replicates commonly used in high-throughput experiments as well as the unique challenges of nucleotide recoding RNA-seq (variability in mutation rates, sequencing errors, heteroskedastic replicate variability, etc.). While there is currently no such tool, several studies provide useful frameworks for interpreting NR-seq data (11, 14, 16). Furthermore, we reasoned that developing such a tool can be guided by numerous advances in the analysis of RNA-seq data to identify differences in RNA levels (i.e., differential expression analysis) (17, 18). In particular, statistical analyses of differential expression are designed to address two important challenges. First, signal is distinguished from background using a statistical model that provides accurate estimates of replicate variability in RNA-seq read counts (19). Second, analyses can compensate for the low number of replicates typically collected for RNA-seq by using transcriptome-wide trends to improve transcript-specific parameter estimates (20–22). Software implementing such analyses (DESeq2 (23), edgeR (24), limma (25), etc.) have become an integral part of almost any RNA-seq experiment, bringing statistical rigor and reproducibility to these high-throughput investigations. New analogous tools are needed for NR-seq data to achieve similar rigor.

To address this need, we developed bakR (<u>B</u>ayesian <u>a</u>nalysis of the <u>k</u>inetics of <u>R</u>NA), an R package that implements an improved statistical model together with multiple-test adjusted null hypothesis testing to perform differential kinetic analyses. bakR consists of three different implementations of a novel Bayesian hierarchical model that we developed to share information across transcripts in high-throughput NR-seq datasets. These implementations allow for both rapid preliminary investigations and more highly powered analyses of differential RNA kinetics. We used a panel of simulated reference and experimental datasets to validate all implementations. We also compared these implementations with implementations we developed that use existing models to perform kinetic parameter estimation prior to differential kinetic analysis (11, 14, 16). We found that all three bakR implementations outperformed similarly efficient implementations of existing models. Finally, we analyzed existing NR-seq datasets with bakR and found that it significantly improved and facilitated the identification of differentially stable transcripts, independent of the NR-seq method employed. bakR represents an important new tool for assessing changes in the kinetics of RNA synthesis and degradation.

## Results

### Adapting existing models and developing a new hierarchical model of NR-seq data

We developed bakR to identify differences in transcript metabolic kinetics by analyzing data from a simple and widely used design for NR-seq experiments (11, 12, 15, 26–32) in which cells are treated with a metabolic label for an amount of time referred to as the label time (t_label_) (Fig. 1A). bakR quantifies how RNA kinetics impacts NR-seq data with a widely-used steady-state model of RNA dynamics (9, 11, 12, 14). In this model, the amount of RNA synthesized during metabolic labeling (RNA_new_) observes first order single rate constant degradation and zeroth order constant rate synthesis kinetics, captured in this differential equation:

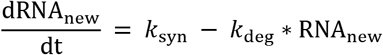

which has the following solution, assuming an initial *RNA*_new_ concentration of zero:

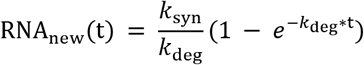

If RNA levels are assumed to be at steady-state, meaning that the total RNA concentration during the experiment is the ratio of *k*_syn_ to *k*_deg_, the fraction new (denoted *θ*) is:

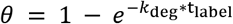

Therefore, to estimate *k*_deg_, *θ* must be estimated. *k*_syn_ can then be estimated using the *k*_deg_ estimate and the total RNA concentration. Estimating *θ* relies on the fact that sequencing reads from new transcripts will have on average more T-to-C mutations (or G-to-A mutations if using s^6^G (33, 34)) than reads from old transcripts.

We considered implementing one of the existing models of NR-seq mutational data to estimate *θ* prior to differential kinetic analysis: pulseR’s complete pooling negative binomial (CPNB) model and a slight modification of Schofield et al. and GRAND-SLAM’s non-hierarchical mixture model (NHPMM) (Fig. 2A; for naming rationale and implementation details see Additional file 2: Supplemental Methods). In addition, we were inspired by models of differential expression to develop implementations of a new hierarchical mixture model (Fig. 2A and B).

**Fig. 2:**
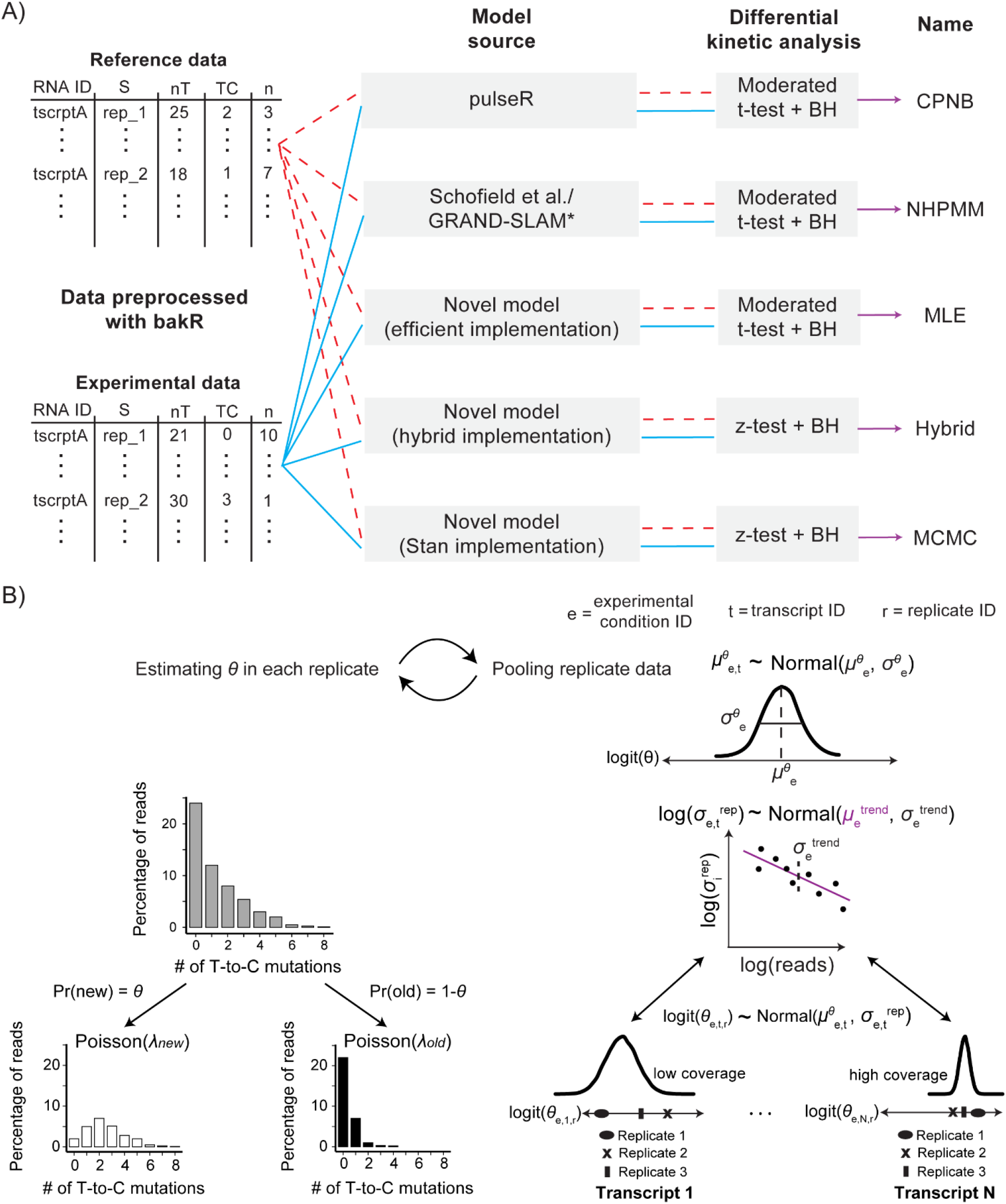
Considering five implementations of new and existing statistical models for bakR. **A)** Names given to the five models/implementations considered for bakR. *Schofield et al./GRAND-SLAM model is a U-content adjusted Poisson mixture model rather than the original binomial mixture model. See **Methods** and Additional file 2: Supplemental Methods for details. **B)** Pictorial description of the new hierarchical mixture model we developed. Mutations in sequencing reads are modeled as coming from a U-content adjusted Poisson distribution two-component mixture model. Partial pooling across fraction new and variance estimates in a given replicate is performed to make use of the high-throughput nature of NR-seq datasets.

Our new hierarchical model was designed to address key limitations of the existing models. The CPNB model assumes that sequencing reads can be unambiguously identified as s^4^U labeled or unlabeled. While this approximation is reasonably accurate in cases of high label incorporation and relatively long reads, it is problematic when faced with lower label incorporation rates and/or shorter (e.g., less than 50 nt) reads (32, 35). Our lab originally addressed this shortcoming by developing a two-component mixture model that accounts for overlap in the mutational distributions of old and new sequencing reads (11) that was implemented by Jurges et al. in the single sample analysis software GRAND-SLAM (14). Despite this improvement, the original mixture model only directly quantifies uncertainty in the fraction new (*θ*) estimate in each replicate and does not attempt to model replicate variability (RV; defined as the standard deviation of the logit(*θ*) estimates, Fig. 3A), an important source of uncertainty in high-throughput, low-replicate number experiments. Thus, we considered strategies to improve uncertainty quantification and directly model mutational data while developing bakR.

**Fig. 3:**
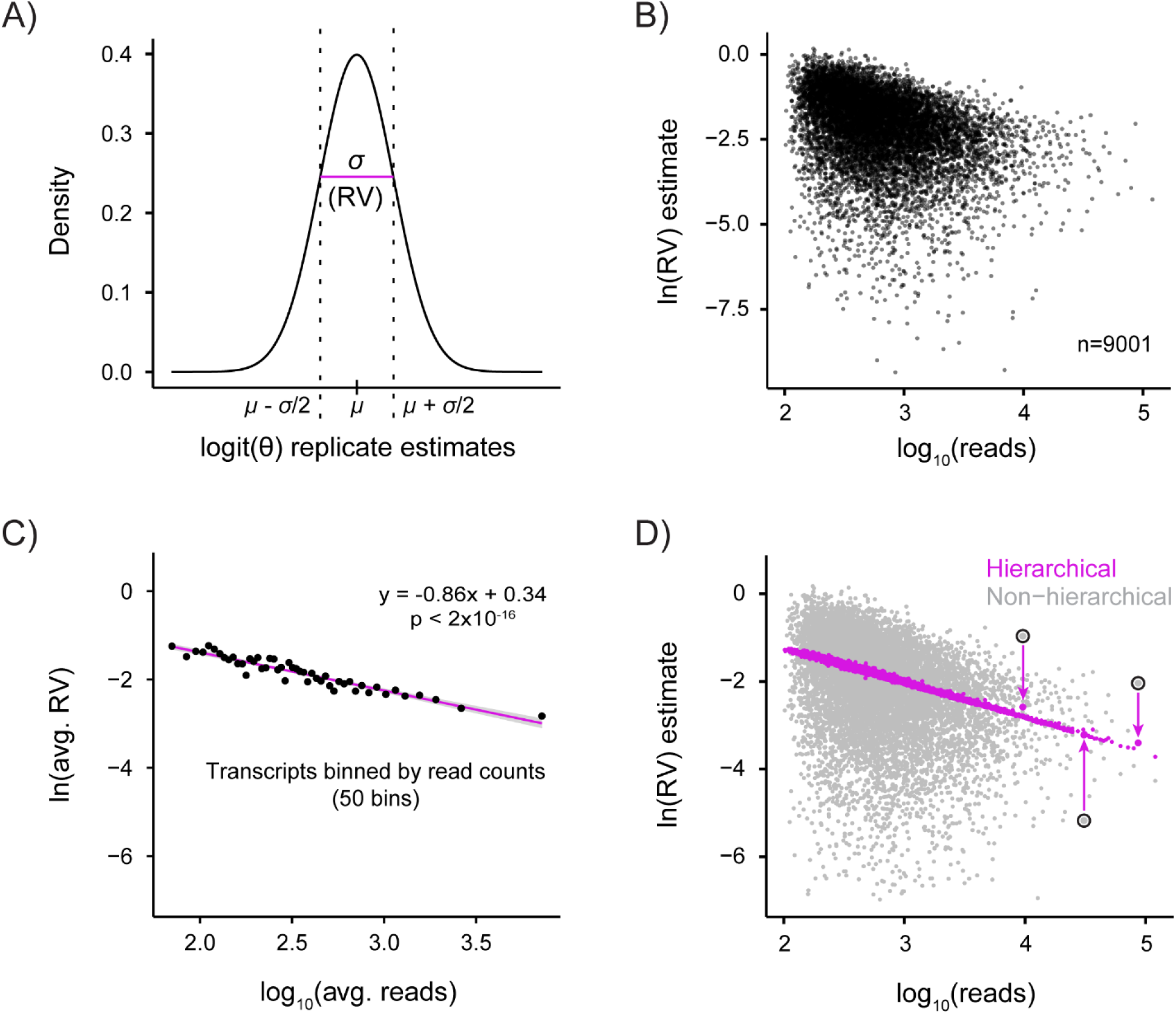
Using a replicate variability trend in NR-seq data to improve uncertainty quantification. **A)** Replicate variability (RV) is defined as the standard deviation of logit(*θ*) estimates across replicates. **B)** RV estimates (using typical sample standard deviation estimator) for individual transcripts in an NR-seq dataset compared to sequencing read counts. **C)** Estimates of average RV for groups of transcripts with similar sequencing depths. **D)** Effect of partial pooling on RV estimate variability. For three transcripts (slightly larger points circled in black), the extent of RV regularization is depicted with an arrow from the naïve estimate to the hierarchical model’s estimate.

NR-seq experiments are typically done with only a few replicates, making it difficult to estimate replicate variabilities for individual transcripts (36). Differential expression analyses for standard RNA-seq have addressed this challenge by sharing replicate variability information between transcripts (23–25). We were particularly inspired by the use of a trend in the replicate variability as a function of sequencing depth that is implemented in limma (25), DSS (37), and DESeq2 (23); therefore, we examined whether there is a similar trend in NR-seq datasets. We found a striking power law relationship between the logit(*θ*) replicate variability (RV) and the number of sequencing reads mapping to a transcript (Fig. 3B, C, and Additional file 1: Fig. S1). In our Bayesian hierarchical model, this trend is used as an informative prior learned from the data that significantly improves RV estimates (Fig. 3D), providing a more accurate separation of signal from background.

### Developing implementations prioritizing either efficiency or accuracy

When considering computational implementations for bakR, we recognized that different applications are likely to favor different needs when balancing statistical rigor/accuracy with computational efficiency. Therefore, we developed three implementations of the novel hierarchical model described above (Fig. 2 and 4A). Efficient maximum likelihood estimation (MLE) combined with partial pooling via analytical solutions to Bayesian models can provide quick but conservative initial investigations of NR-seq datasets. A more sensitive but less efficient full Markov Chain Monte Carlo (MCMC) implementation using the probabilistic programming language Stan (38) can provide a more comprehensive statistical analysis. Finally, we reasoned that a hybrid approach (Hybrid implementation) that combines features of the MLE and MCMC implementations would be particularly useful when analyzing large datasets (e.g., 20 samples or more each with at least 10 million mapped reads), as a full MCMC implementation might take anywhere from weeks to months to run on such datasets. See **Methods** and Additional file 2: Supplemental Methods for further details regarding all implementations. The CPNB implementation is of similar efficiency to that of the MLE implementation and the NHPMM implementation is of similar efficiency to that of the MCMC implementation (Additional file 1: Table S1). We next examined the performance of these potential bakR implementations.

**Fig. 4:**
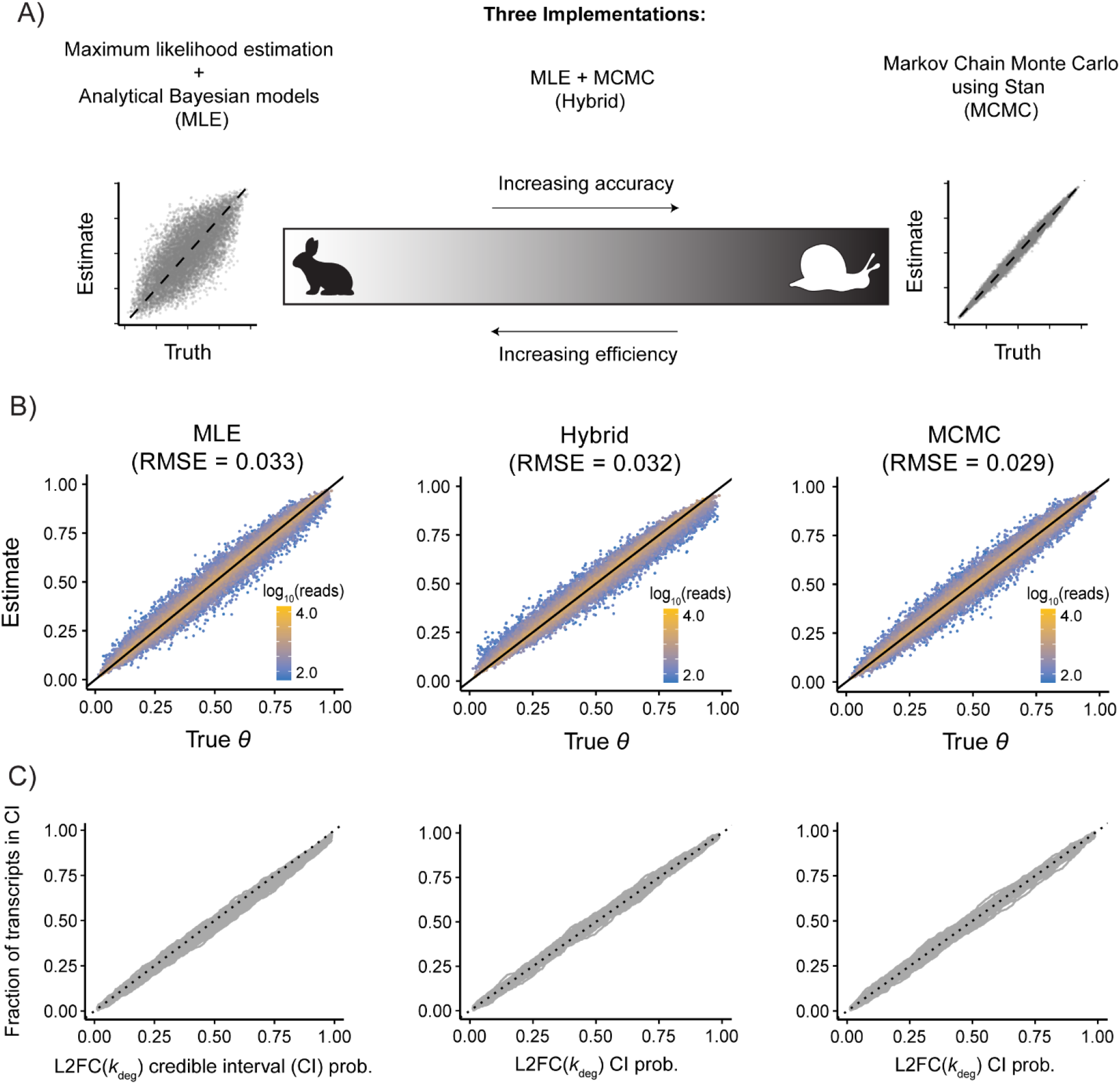
Development and validation of three distinct computational implementations of the Bayesian hierarchical model. **A)** Fundamental tradeoff exists between efficiency and accuracy of a statistical model’s computational implementation. bakR employs three distinct computational implementations that cover the efficiency vs. accuracy spectrum. **B)** Accuracy of *θ* estimates for the three implementations of the new hierarchical model. Diagonal line is y = x. **C)** L2FC(*k*_deg_) calibration for all three implementations of the new hierarchical model. The x-axis represents the marginal posterior probability mass covered by the L2FC(*k*_deg_) credible interval and the y-axis is the percentage of transcripts who’s true L2FC(*k*_deg_) fall within the credible interval. 500 lines obtained from bootstrapped samples of 1000 transcripts from a total of 10000 transcripts are depicted for each implementation.

### Validation of all five implementations using simulated data

We used simulated data to assess the accuracy of all implementations considered for bakR (see **Methods** for simulation details). Briefly, we used a generative model that captures trends seen in real NR-seq datasets. We simulated two replicates of NR-seq data for 10,000 transcripts and two experimental conditions (e.g., control and treated samples). We analyzed the simulated data with all five implementations. This analysis revealed strong correlations between the estimates and the true simulated values (Fig. 4B, and Additional file 1: Fig. S2), confirming that *θ* estimates from the three implementations of the new hierarchical model were accurate. *θ* estimate accuracy was found to be greater for simulated transcripts with more reads, as expected. The CPNB and NHPMM estimates were also generally accurate, though the CPNB estimates were worse for transcripts with *θ* further from 0.5 (Additional file 1: Fig. S3). This is likely due to pulseR’s assumption that labeled and unlabeled reads can be definitively identified. In summary, *θ* was well estimated by all implementations, especially those directly modeling the mutational data.

As bakR is the first NR-seq analysis tool to perform differential kinetic analysis, we next confirmed that estimates of log2-fold changes in the degradation rate constant (L2FC(*k*_deg_)) were accurate and that uncertainties were properly quantified by all potential bakR implementations. To do this, we assessed the extent to which the credible/confidence intervals provided by the five implementations contained the true L2FC(*k*_deg_) with the expected probability (Fig. 4C and Additional file 1: Fig. S3). For example, the 50% credible interval should contain the simulated L2FC(*k*_deg_) 50% of the time, and we found this to be generally true for all implementations.

Finally, a benefit of Stan’s implementation of auxiliary variable MCMC, used by the NPHMM, Hybrid, and MCMC implementations, is that it provides robust model diagnostics (39, 40). These diagnostics can be used to confirm that all model parameters are accurately estimated with the data provided. When analyzing simulated data, these sampling diagnostics indicated that this was the case (Additional file 1: Fig. S4 and S5), further supporting the conclusion that the models are well implemented.

### Simulated data comparison of models and implementations

We next determined which of the five models/implementations to include in bakR by comparing their ability to distinguish differential stability signal from null background in multiple simulated datasets. We simulated data for 10,000 transcripts, of which 10% were simulated to have differential degradation kinetics (differences in stability between a reference and experimental condition; simulated L2FC(*k*_deg_) magnitudes typically ranged from just above 0 to just under 3), two biological replicates of data, and three different read count vs. replicate variability trends (Fig. 5 top row; see **Methods** for simulation details). These replicate variability trends represent the range of trends we have seen in real NR-seq datasets. The false discovery rate (FDR, percentage of transcripts identified by the models as hits that were false positives) and power (percentage of transcripts with differential kinetics identified as hits by the models) of all models were assessed at a range of significance cutoffs (Fig. 5 middle and bottom rows, respectively). FDR was calculated at a wider range of cutoffs to assess the extent to which FDR was controlled, as the ideal case is a *y* = *x* relationship between p_adj_ cutoff and FDR. Power was calculated for a range of commonly used significance cutoffs (p_adj_ < 0.01 to p_adj_ < 0.1). We also compared Matthews correlation coefficients (MCC) (41) of all the models at a common significance cutoff of 0.05 (Additional file 1: Fig. S6). The MCC is more informative than other popular binary classification metrics (e.g., F_1_ score and model accuracy) as it accounts for the relative abundances of all four categories of classifications (i.e., false positives, false negatives, true positives, and true negatives) (42).

**Fig. 5:**
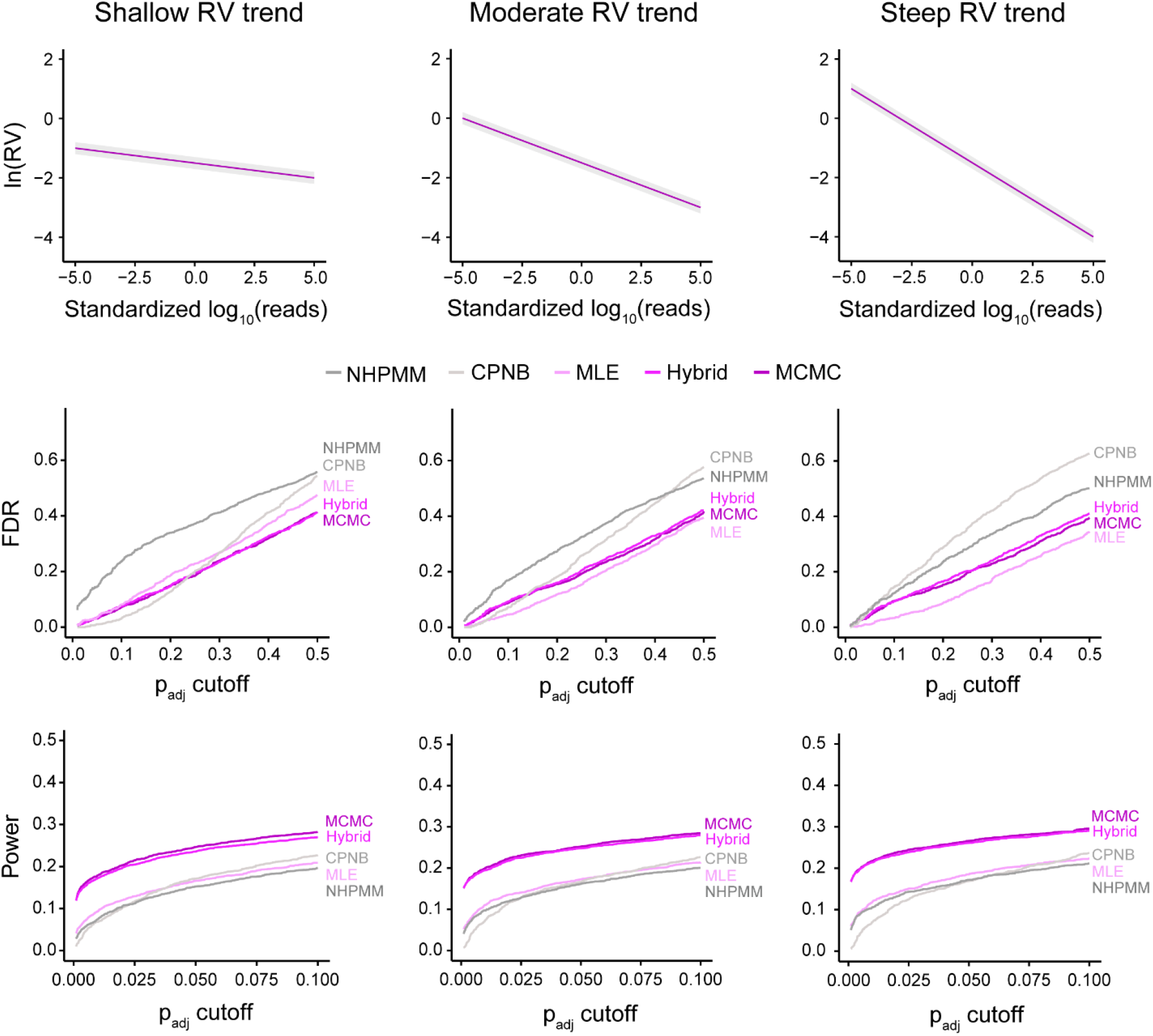
bakR implementations outperform extensions of existing models. **Top:** Simulated RV vs. read count trend. ln(RV) for each transcript is drawn from a normal distribution centered on the trend with standard deviation of 0.05. Gray bar around trend represents typical range of simulated replicate variabilities. **Middle:** False discovery rate (FDR) as a function of significance cutoff for all implementations. **Bottom:** Power as a function of significance cutoff for all implementations. The average FDR and Power for 5 simulation replicates are plotted at each cutoff.

In all cases, we found that FDRs were well controlled by the Hybrid and MCMC implementations (Fig. 5 and Additional file 1: Fig. S7). FDRs for the NPHMM implementation were slightly inflated. Despite this, the NPHMM implementation was still significantly underpowered relative to the Hybrid and MCMC implementations. In other words, the Hybrid and MCMC implementations identified more signal and filtered out more null transcripts than the NPHMM implementation. Therefore, we decided that the Hybrid and MCMC implementations were optimal for analyses of NR-seq datasets.

Comparing the more efficient implementations (MLE and CPNB), FDRs were generally well controlled in both cases, though became more inflated for the CPNB implementation when analyzing simulated data with steeper read count vs. replicate variability trends (Fig. 5). The statistical power and MCCs of these implementations were also comparable. Increasing the amount of variability in the replicate variability, decreasing s^4^U incorporation rates, and increasing the number of replicates did not significantly alter the relative performance of these implementations (Fig. 5 and Additional file 1: Fig. S7). Due to its improved kinetic parameter estimation accuracy (Fig. 4B and Additional file 1: Fig. S3), robustness to different heteroskedastic trends (Fig. 5), superior efficiency (Additional File 1: Table S1), and compatibility with the Hybrid implementation, we included the MLE implementation and not the CPNB implementation in bakR. Thus, bakR includes the MLE, Hybrid, and MCMC implementations.

### bakR improves and facilitates analyses of real NR-seq datasets

We next examined the performance of bakR using experimental data. We first reanalyzed Luo et al.’s TimeLapse-seq data with RNA collected from WT and *DCP2* knock-out (KO) HEK293T cells (15). The DCP2 enzyme catalyzes the removal of the 5′ protective cap of mRNA (43), so the expectation is that *DCP2* KO will lead to widespread stabilization of DCP2 mRNA substrates. Lacking a tool like bakR, Luo et al. developed an analysis strategy specialized for their data. This strategy was able to accurately quantify changes in degradation rate constants but lacked uncertainty quantification and thus relied on differential expression analysis and comparison of L2FC(*k*_deg_) vs. L2FC(*k*_syn_) magnitudes to identify stabilization (see **Methods** for details). Therefore, we hypothesized that bakR could significantly improve the list of transcripts identified as stabilized in *DCP2* KO cells.

Reanalysis of the data from Luo et al. using all three implementations of bakR confirmed that there was widespread stabilization of transcripts in the *DCP2* KO (Fig. 6B and Additional file 1: Fig. S8; FDR < 0.05). The MCMC and Hybrid implementations both identified over 1700 stabilized transcripts, and the MLE implementation identified 529 stabilized transcripts. While the number of transcripts identified as stabilized by the previous analysis strategy (1692) was comparable to bakR’s MCMC implementation, we noticed that there were significant differences in the identity of these transcripts (Additional file 1: Fig. S9A). The differences were not driven by differences in L2FC(*k*_deg_) estimates (Additional file 1: Fig. S9B), which were quite similar across analyses. Rather, there were two major causes of the discrepancy. First, the previous analysis did not quantify the uncertainty in the L2FC(*k*_deg_) estimate and in many cases inferred stabilization of low coverage transcripts with insufficient data (Fig. 6C and Additional file 1: Fig. S9C and Fig. S10). Second, the previous analysis only considered differentially expressed transcripts, causing it to miss many instances of highly stabilized transcripts that were not identified as significantly upregulated (Additional file 1: Fig. S9D, E, and F and Fig. S10). Relying on differential expression analysis can be limiting especially in a case like this where widespread stabilization is expected and the experiment was performed without spike-ins for read count normalization (44). Alternatively, even with additional normalization, transcripts with significant stabilization in absence of upregulation could be due to the increasingly recognized phenomenon of compensatory regulation of transcription to buffer changes in transcript stability (45–50). Either way, bakR provides the first list of *DCP2* KO stabilized transcripts that accounts for L2FC(*k*_deg_) estimate uncertainties.

**Fig. 6:**
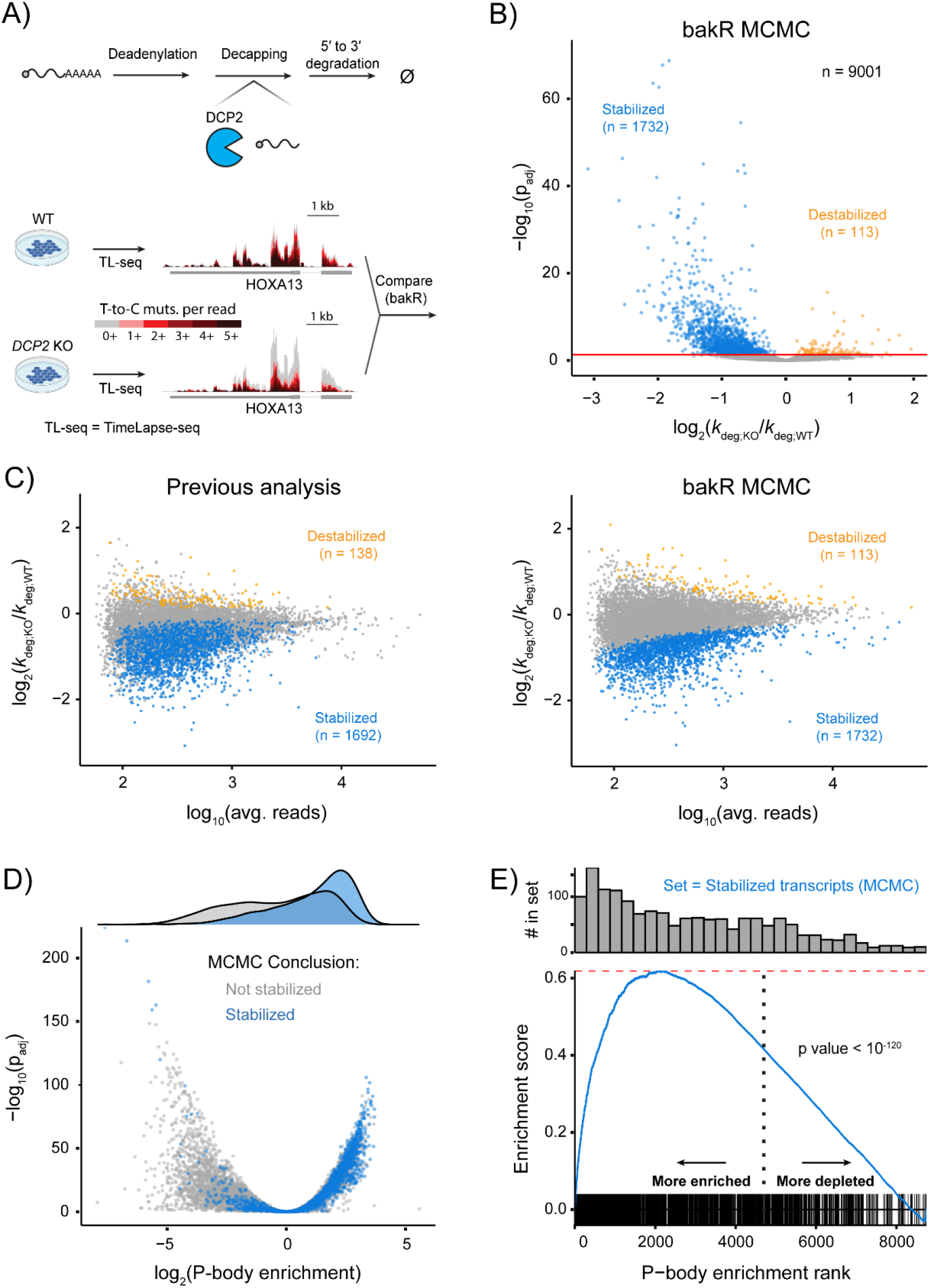
bakR provides an improved statistical foundation to identify biological signals in an existing NR-seq dataset. **A)** Experimental setup used by Luo et al. Transcript stabilities in WT HEK293T were compared to those in *DCP2* knockout (KO) cells. **B)** Volcano plot obtained from analysis of data with bakR’s MCMC implementation. **C)** Comparison of MA plots obtained with the previous analysis strategy of Luo et al. and bakR’s MCMC implementation **D)** Correlation between P-body enrichment and DCP2-dependent degradation. **E)** GSEA analysis using the MCMC stabilized transcripts as the gene set of interest and P-body enrichment scores as the scoring function.

We next examined the biological implications of the improved list. Luo et al. previously found that transcripts categorized as stabilized in the *DCP2* KO are often associated with P-bodies, which are phase separated granules where the DCP2 protein localizes (51). We sought to reexamine P-body enrichment while showcasing a promising application of bakR by performing gene-set enrichment analysis (GSEA) using the *DCP2* KO-stabilized transcripts derived from bakR (FDR < 0.05) and P-body enrichment scores from a prior study (52). As expected, we found that stabilized transcripts were significantly enriched among P-body transcripts (Fig. 6D, E, and Additional file 1: Fig. S8). Furthermore, bakR amplified this biological signal relative to the previous analysis strategy (Additional file 1: Fig. S11). These results confirm that bakR provides an improved statistical foundation to identify biological signals in NR-seq datasets.

To test the generality of bakR using other NR-seq methods, we reanalyzed data from SLAM-seq with total RNA from *Mettl3* KO mouse embryonic stem cells (mESC) and WT mESC (12). Mettl3 is a writer of the m^6^A RNA modification, a mark which has been found to promote degradation of modified transcripts (53). Using bakR, we identified widespread stabilization in the *Mettl3* KO (Additional file 1: Fig. S12) consistent with previous studies. In addition, we performed GSEA analysis but this time using bakR’s L2FC(*k*_deg_) z-score as the scoring function and a list of previously identified Mettl3 targets as the gene set of interest (54). Mettl3 targets were significantly enriched among the most stabilized transcripts (Additional file 1: Fig. S12). These analyses require the uncertainty quantification provided by bakR and confirm that bakR effectively identifies biological signal while also being applicable to different NR-seq protocols.

## Discussion

bakR is the first tool developed to perform differential kinetic analysis using statistical modeling of nucleotide recoding RNA-seq (NR-seq) data. While existing software can analyze individual NR-seq samples (14, 16), bakR performs the necessary statistical inference to compare NR-seq datasets. In addition, bakR implements a new Bayesian hierarchical model that improves uncertainty quantification and statistical power in these typically low replicate number, high-throughput datasets.

We included three distinct computational implementations of our Bayesian hierarchical mixture model in bakR (MLE, Hybrid, and MCMC). While the MCMC and Hybrid implementations significantly outperform all other tested implementations, their longer runtimes often necessitate the use of specialized computation infrastructure (e.g., High Performance Computing) when analyzing typical NR-seq datasets. Therefore, we also provide the MLE implementation as an efficient and conservative alternative to facilitate rapid exploratory investigations of differential RNA metabolic kinetics.

We were struck by the comparable performance of the Hybrid and MCMC implementations in most of our simulations (Fig. 5 and Additional file 1: Fig. S6 and S7). These implementations often had a similar FDR, power, and MCC at all significance cutoffs assessed. Despite this as well as the Hybrid implementation’s superior efficiency, we included the MCMC implementation in bakR because it offers advantages over the other implementations. The MCMC implementation provides mutation rate estimates that are typically more robust than those provided by the MLE implementation, especially in difficult experimental settings (e.g., short sequencing reds, low mutation rates, etc.; Additional file 1: Fig. S7 for example). The MCMC implementation also includes sampling diagnostics provided by Stan (38). Failed convergence of the MCMC implementation’s Stan model may indicate subtle anomalies in NR-seq datasets that the other implementations and bakR quality control functions cannot identify. The fully Bayesian model fit provided by the MCMC implementation also opens the door for additional explorations of NR-seq datasets (e.g., mutational data posterior predictive checks (39)). Thus, all three computational implementations provided by bakR offer complementary advantages.

Analysis of simulated data confirmed that bakR can accurately estimate kinetic parameters and differences in kinetic parameters between two samples. Simulations also demonstrated that bakR significantly outperforms differential kinetic analysis with extensions of existing models (the CPNB and NHPMM implementations). We analyzed several existing NR-seq datasets (TimeLapse-seq data in *DCP2* KO and WT 293T cells (15) and SLAM-seq data in *Mettl3* KO and WT mESCs (12)) and found that bakR can effectively identify differences that capture biologically relevant changes.

bakR paired with metabolic labeling, nucleotide recoding chemistry, and RNA-seq significantly facilitates statistically rigorous differential RNA kinetic analysis. bakR uses Bayesian hierarchical modeling to effectively share information across transcripts, increasing its statistical power while maintaining the desired level of specificity. We anticipate that broad implementation of bakR to analyze NR-seq datasets will provide new insights into the regulated kinetics of gene expression.

### Limitations

bakR implements a Bayesian hierarchical model and downstream statistical inference to facilitate comparisons of RNA kinetics in two or more samples. Currently, bakR is only compatible with comparisons of a set of experimental samples to a single reference sample. Additional comparisons can be performed with bakR, but multiple test adjustment that accounts for the true number of hypothesis tests conducted must be implemented manually.

In addition, bakR does not perform differential expression analysis, so the output of well-established differential expression software like DESeq2 must be combined with the output of bakR to perform differential RNA synthesis kinetic analysis. We chose to avoid redundant development of differential expression analyses in bakR since several well-validated tools exist for this purpose.

bakR relies on metabolic labeling and nucleotide recoding chemistry, a powerful experimental approach for studying gene expression kinetics (9), yet not all biological systems are compatible with metabolic labeling. In those cases, other methods can be used to extract kinetic information from RNA-seq data without metabolic labeling by modeling intronic and exonic read counts (3, 55). Unfortunately, these models are underdetermined unless provided with multiple time points of RNA-seq data. Therefore, when possible, NR-seq combined with bakR provides more statistical power to assess differential RNA kinetics with fewer samples.

## Conclusions

In this work we presented bakR, the first tool designed to estimate and compare RNA synthesis and degradation kinetic parameters using NR-seq. Using simulated and real datasets, we showed that bakR’s Bayesian hierarchical modeling significantly improves our ability to identify differences in RNA kinetics. Of the numerous publications using NR-seq to date, the vast majority have applied NR-seq to the task of differential kinetic analysis. Therefore, bakR represents a much-needed tool in this growing field.

## Methods

### Estimation of the fraction new (*θ*)

Both of the mixture models considered (new hierarchical model and NHPMM model) were similar to that described by Schofield et al. (11) and implemented in GRAND-SLAM (14). Briefly, we model the number of mutations in sequencing reads from a given transcript (subscript t) in a given sample (subscript s) as coming from a mixture of two Poisson distributions. One component of the mixture represents the background mutation rate due to sequencing errors in old RNA while the other component models the higher s^4^U recoding augmented mutation rate in new RNA. The likelihood function for this model is:

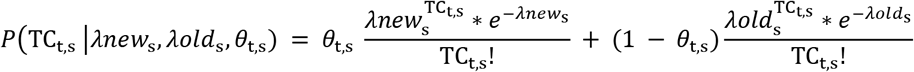

where TC_t,s_ is the number of T-to-C mutations in a single sequencing read, *λnew* is the mutation rate in new RNA, and *λold* is the mutation rate in old RNA. The original models used a binomial mixture model rather than a Poisson mixture model. A Poisson model yields significant compression of input data and far greater efficiency than a binomial model because it does not track the number of Us in each sequencing read. *λnew* and *λold* are global parameters shared by all transcripts, so we account for differences in transcript U-contents by calculating the fold difference in transcript specific and global U-content as:

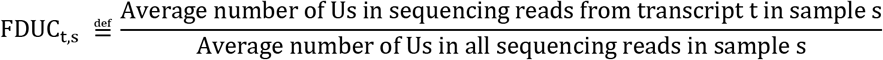

making the U-content adjusted likelihood for transcript t in sample s:

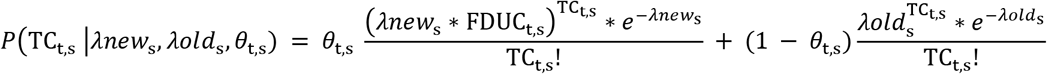

We used simulated data to confirm that the U-content adjusted Poisson mixture model performs as well as a binomial mixture model (Additional file 1: Fig. S13).

### Bayesian hierarchical model

Existing implementations of the mixture model discussed above do not take full advantage of the high-throughput nature of nucleotide recoding RNA-seq datasets. In such datasets, hierarchical models are often optimal (21). Hierarchical modeling uses trends in similar groups of transcripts to improve parameter estimates for individual transcripts (56). In bakR, *θ* and RV (*σ*^rep^) estimates from the same sample are partially pooled to achieve this regularization (Fig. 2). To address the challenges inherent to the implementation of hierarchical models, bakR uses either analytical solutions to simple Bayesian models or Hamiltonian Monte Carlo (HMC) implemented in Stan (38) to sample from the posterior. As discussed in **Developing implementations prioritizing either efficiency or accuracy**, we developed several implementations of our Bayesian hierarchical model for use in bakR. The following hierarchical generative model is used by all implementations:

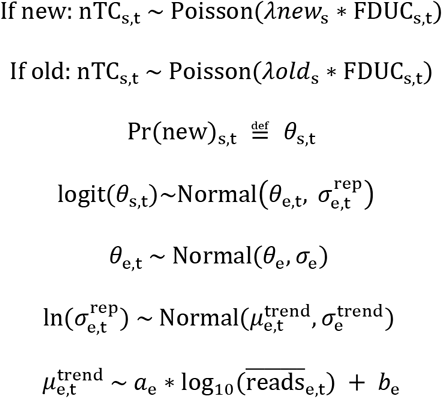

nTC_s,t_ is the number of mutations (T to C mutations if using s^4^U) in a sequencing read, the subscript s is a numerical identifier for each s^4^U treated sample, and the subscript t is a numerical identifier for each transcript analyzed. *λnew* is the s^4^U induced mutation rate, estimated on a sample-wide basis to improve mutation rate estimation. *θ*_e,t_ is the average *θ* for a transcript, averaged over all samples (i.e., replicates) in the same experimental condition, with numerical identifier e. For example, in the *DCP2* KO data reanalyzed in this paper, there are two experimental conditions: wild-type (WT) and the *DCP2* KO (KO). 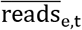 is the average number of reads mapping to transcript t in experimental condition e. Improved model convergence was achieved through passing the model standardized 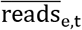 (i.e., subtracting the mean and dividing by the standard deviation for all 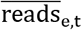 in experimental condition e). 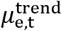 captures where a given transcript falls along the estimated RV vs. read count trend and is used as an informative prior to regularize RV estimates (Fig. 3C and D). The average *θ* (*θ*_e,t_) is estimated for each of the conditions. Partial pooling of *θ* is not only done across replicate *θ* estimates for each transcript (*θ*_s,t_); the average *θ* for all transcripts are also partially pooled. In the MCMC and Hybrid implementations, this avoids the problem of having to explicitly specify a *θ*_e,t_ prior for each transcript.

### MCMC implementation priors and parameterization

This section discusses the specifics of the parameterization and prior for the Stan model used in the MCMC implementations. The specifics of the other two bakR implementations (MLE and Hybrid) are discussed in the Additional file 2: Supplemental Methods. Details about the other two implementations tested (CPNB and NHPMM) are also discussed in the Additional file 2: Supplemental Methods.

When implementing the generative model presented in **Partial pooling of** *θ* **estimates**, there are multiple mathematically equivalent parameterizations that differ in their computational performance (e.g., centered and non-centered parameterizations) (56). Centered parameterizations are typically optimal if the number of parameters being partially pooled is large, while a non-centered parameterization is typically optimal if the number of parameters being partially pooled is small (57). The parameterizations chosen for the MCMC implementation (and the Hybrid implementation, see Additional file 2: Supplemental Methods) largely follow this trend. In contrast, we found that 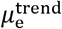, which pools replicate variability data across all transcripts in a sample, is better non-centered, perhaps due to features of hierarchical linear models (39).

The priors for all parameters not modeled as being drawn from distributions of other learned parameters are as follows:

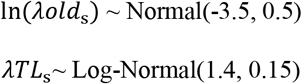

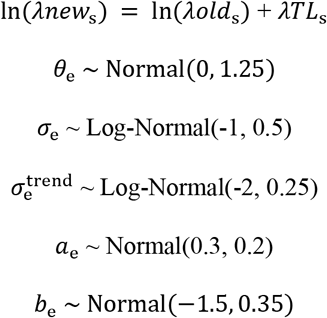

Note, rather than having the model estimate the new and old read mutation rates independently, the model estimates the old mutation rate as well as a strictly positive parameter (*λTL*_e_) to avoid challenges from label-switching (58). Prior distribution parameters were chosen according to trends seen in real data as well as the use of prior predictive distributions. Prior predictive distributions reveal that the prior fraction new distribution is symmetric about 0.5 (or 0 on logit-scale) and covers a reasonable range of values (Additional file 1: Fig. S14A and B). Similarly, prior predictive distributions for a theoretical transcript with an average *θ* of 0.5 shows that the priors capture the increased replicate variability expected for lower sequencing depth features, without predicting extreme amounts of replicate variability (or lack thereof; Additional file 1: Fig. S14C and D).

The average degradation rate constants for each transcript and the log2-fold change in the degradation rate constant (comparing the user defined reference sample to each of the experimental samples) are calculated from the data and estimated parameters as follows:

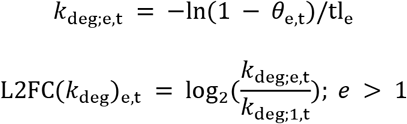

where *k*_*deg*;1,*t*_ is the average degradation rate constant estimate in the reference sample (e = 1). The L2FC(*k*_*deg*_)_*e*,*t*_ marginal posterior distribution is central to all statistical testing performed by bakR, as discussed in the next section

### Differential kinetic analysis

The MCMC, Hybrid, and MLE implementations of our novel hierarchical model provide marginal posterior distributions for the L2FC(*k*_deg_), comparing any number of samples to a reference sample. To compare the binary classification performance of these implementations we used a frequentist multiple-test adjusted null hypothesis testing strategy. First, a test-statistic (denoted *Z*) is calculated using the L2FC(*k*_deg_) marginal posterior mean and standard deviation (sd):

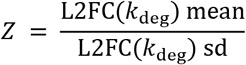

which is then used to calculate a p value using a z-test (i.e., comparing to a normal distribution) for the MCMC and Hybrid implementations and a moderated t-test developed for multi-level linear models for the MLE implementation (59). The p value is then multiple-test adjusted using the method of Benjamini and Hochberg (60). We initially tried using a moderated t-test for all implementations but found it to be consistently overconservative with the MCMC and Hybrid implementations, likely due to the additional partial pooling of these models. For these implementations, the z-test was found to yield statistical conclusions that more accurately reflected the FDR cutoff used (see the agreement between p_adj_ cutoff and FDR for the MCMC and Hybrid implementations in Fig. 5 for example).

The CPNB model (pulseR) provides a single estimate averaged across replicates of a particular biological condition, as well as a confidence interval estimated using a profile likelihood approach (16). Thus, CPNB takes pulseR’s degradation rate constant estimates from two different conditions and calculates the log2-fold difference in each transcript’s *k*_deg_. To estimate the standard error of the L2FC(*k*_deg_), we assumed independence of the two estimates, in which case the standard error is the square root of the sum of the squares of the standard errors of the two log_2_(*k*_deg_) estimates. Standard errors of the log_2_(*k*_deg_) estimates were approximated as one-quarter of the width of the 95% confidence intervals. We used the L2FC(*k*_deg_) estimate and standard error to calculate a t-statistic for each transcript, that being the ratio of these two quantities. We used the same two-tailed moderated t-test as for the MLE implementation. The p value was subsequently adjusted for multiple testing using the Benjamini-Hochberg method. The NHPMM model implementation provides a log_2_(*k*_deg_) estimate for each transcript in each replicate of a particular biological condition. We first pooled replicate data to determine a condition-wide average, and then performed the same downstream analysis as described for the CPNB estimates. The condition-wide average was calculated as the weighted mean of the replicate estimates, weighted by the inverse of the square of the posterior standard deviation. The condition-wide standard error was calculated as:

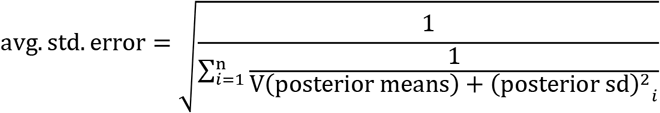

where V(posterior means) is the sample variance of the posterior means across the n replicates. Statistical testing was performed as with the CPNB implementation.

### Simulations for model validation and comparison

We used simulated data to validate bakR’s three implementations and to compare these implementations to extensions of pulseR and the model from Schofield et al/GRAND-SLAM. To eliminate dependence on upstream processing of fastq files, we chose to directly simulate data in a form that could be analyzed by existing statistical models. This was done with a generative model that captures trends seen in real data. The model consists of two sub-models: a model for the mutational content of sequencing reads in a single replicate and a model for the variability across replicates. A mathematical description of the model used to simulate data for all transcripts in all replicates of the reference condition is presented below; for the experimental condition the only difference is that a L2FC(*k*_deg_) is simulated and combined with the reference θ_*o*_ to simulate experimental condition kinetic parameters. A single replicate of an unlabeled control is also simulated for both experimental conditions, in which case the only difference is that all mutations are drawn from the binomial distribution used for old reads in labeled samples:

#### Explanation of indices used

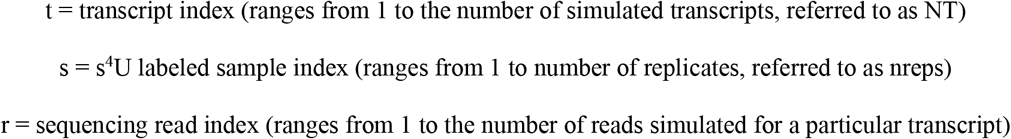

#### Simulating mutational data in reads

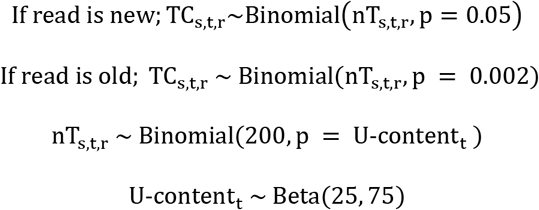

#### Simulating kinetic parameters and replicate variability for reference sample

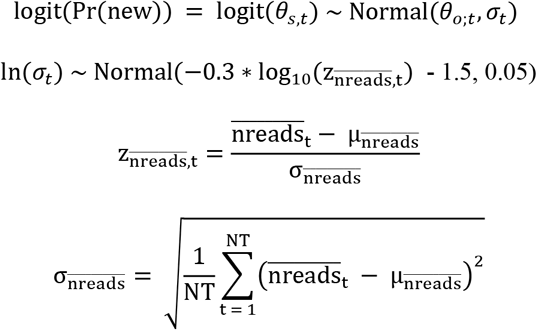

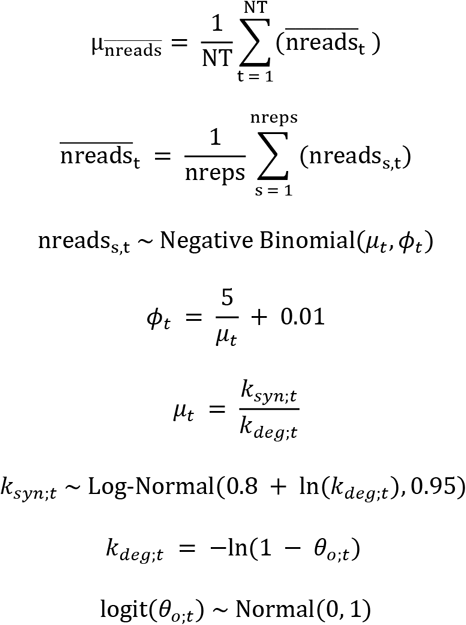

Two conditions were always simulated (one reference and one experimental), and the number of replicates simulated was either 2 or 3. For the experimental condition, 90% of transcripts were chosen to have 0 average difference in stability between the two conditions. The rest of the transcripts had a L2FC(*k*_deg_) drawn from a Normal distribution with mean 0 and standard deviation 0.75. This distribution of L2FC(*k*_deg_)s captures the fact that for many experimental treatments there are a range of effect sizes, and is similar to effect size distributions put used elsewhere (61). A typical range for |L2FC(*k*_deg_)| is from 1e-4 to 2.6, which is reflective of that seen in the NR-seq datasets we analyzed in this paper. All other parameters were chosen so that the simulated data matched trends seen in WT HEK293T TimeLapse-seq data. This simulation strategy is implemented in bakR as the Simulate_bakRData function.

### Pipeline to preprocess existing TimeLapse-seq and SLAM-seq datasets

Filtering and alignment to the human GRCh38 genome release 104 (for TimeLapse-seq data) were performed essentially as described previously with some modifications (11). Briefly, reads were filtered for unique sequences using FastUniq v.1.1 (62) and trimmed of adaptor sequences with Cutadapt v3.2 (63). Reads were aligned with HISAT-3N (64) with default parameters. Reads aligning to annotated transcripts were quantified with HTSeq (65) htseq-count. SAMtools v.1.5 (66) was used to collect only uniquely mapped read pairs (SAM flag = 83/163 or 99/147). Mutation calling was also performed essentially as described previously. Briefly, T-to-C mutations were not considered if the base quality score was less than 40. In addition, mutations within 5 nucleotides of the read’s end were not considered. Sites of likely single-nucleotide polymorphisms (SNPs) and alignment artefacts (identified with bcftools v.1.11 (67)) were not considered in mutation calling. Multi-thread parallelization was implemented with GNU parallel 20210222 (68). Filtering and alignment of SLAM-seq 3′ end sequencing data to the mouse GRCm39 genome release 105 was performed almost identically with some modifications. Reads were single end so FastUniq was not used, and a regular 5′ adapter of AAAAAAAA was provided to Cutadapt in addition to the regular 3′ adapter of AGATCGGAAGAGC to trim the polyA tail.

### Analysis of existing TimeLapse-seq data

bakR’s default filtering criteria for all implementations includes removing transcripts with less than 50 mapped reads in any sample and removing transcripts with abnormally high mutation rates (> 20%) in any unlabeled control samples. The filtered data was then analyzed with all three implementations using default settings except for mutation rate estimation for the MLE implementation. In that case, StanRateEst was set to TRUE in the bakRFit function to improve mutation rate estimation in this implementation. See Additional file 2: Supplemental Methods for details.

To compare the bakR analysis with previous analyses, we reanalyzed the TimeLapse-seq data using the strategy described by Luo et al. Briefly, we restricted comparisons to transcripts identified as upregulated by DESeq2 (L2FC > 0, p value < 0.05), and estimated degradation rate constants using a CPNB-like assumption that reads can be separated as new and old based on their mutation rate. Simulations had previously determined an optimal new versus old cutoff (> 3.6% of Us mutated to Cs) at the observed mutation rates. Thus, the fraction new (*θ*), degradation rate constant (*k*_deg_), and *k*_deg_ log2-fold change (L2FC(*k*_deg_)) were estimated as:

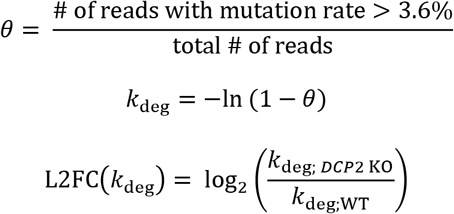

The L2FC(*k*_deg_) and DESeq2’s L2FC([*RNA*]) estimates were combined to estimate the L2FC(*k*_syn_):

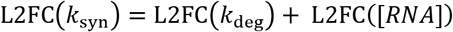

Finally, stabilized transcripts were identified as those for which the magnitude of the L2FC(*k*_deg_) was greater than the magnitude of the L2FC(*k*_syn_).

For the GSEA analysis, we used the fgsea R package (69). The score for each transcript was the −log_10_ transformed p value for the P-body enrichment of that transcript, multiplied by the sign of the enrichment (i.e., multiplied by 1 if the transcript was enriched in P-bodies and −1 if the transcript was depleted from P-bodies). Transcripts with a p value of 0 were assigned the maximum absolute score plus a Uniform(0,1) jitter, and multiplied by the sign of the enrichment. In Additional file 1: Fig. S11, we applied an additional log10 transformation to the P-body enrichment score (i.e., |log_10_| of the −log_10_ p value, again multiplied by the sign of the enrichment). We used the fgsea function with eps set to 0 to assess enrichment of a single gene set, genes from which transcripts identified as stabilized by bakR are synthesized.

### Analysis of existing SLAM-seq data

Filtering and analysis with bakR were performed as described for the reanalysis of an existing TimeLapse-seq dataset. GSEA was performed similarly as with the TimeLapse-seq data, but with the following modifications. The gene set for which enrichment was assessed were genes from which transcripts previously identified as having Mettl3 targets in their coding sequence (CDS) were synthesized. The GSEA scoring function was the L2FC(*k*_deg_) z-score from bakR. We note that bakR improves on the previous analysis strategy employed on this dataset in two ways (12). One, the previous analysis required the use of data from two separate s^4^U label times (3 hours and 12 hours), whereas bakR was able to obtain quality estimates from a single label time (3 hours). Secondly, the previous analysis lacked L2FC(*k*_deg_) uncertainty quantification and thus was not able to provide an ordered list of *Mettl3* KO-stabilized transcripts.

## Declarations

### Availability of data and materials

bakR is freely available under an MIT license on Github (https://github.com/simonlabcode/bakR). Data and scripts required to reproduce all main text and supplemental figures are available on OSF (link: https://osf.io/4eug7/?view_only=ec68db829c7b4d82a81ffd9fcc076ffb, DOI: 10.17605/OSF.IO/4EUG7). Analyses presented in this paper were performed with bakR version 0.1.0. The pipeline implementing the fastq processing described in **Pipeline to preprocess existing TimeLapse-seq and SLAM-seq datasets** is available on bitbucket (https://bitbucket.org/mattsimon9/timelapse_pipeline/src/master/). We used the following publicly available datasets:

- TimeLapse-seq data in WT and *DCP2* KO HEK293T cells (15)
  - GEO accession number GSE143662
- SLAM-seq data in WT and *Mettl3* KO mESC (12)
  - GEO accession number GSE99978

### Authors’ contributions

MDS conceived the idea and provided funding support. IWV developed the Bayesian hierarchical model and implemented the software. IWV performed simulations and analyzed real data. IWV and MDS wrote the manuscript.

### Consent for publication

Not applicable

### Ethics approval and consent to participate

Not applicable

## Acknowledgements

The authors thank all members of the Simon lab for useful discussions during the building and testing of bakR. In addition, we want to thank Martin Machyna and Joshua Zimmer for improvements to computational analysis of NR-seq data. We thank Joshua Zimmer and Andreas Pintado-Urbanc for helping with beta-testing and providing thoughtful feedback, as well as Michelle Moon for useful suggestions regarding bakR’s input object. Finally, we want to thank Wendy Gilbert, Lea Kiefer, Joshua Zimmer, and Michelle Moon for helpful comments on this manuscript.

## References

1. Garneau NL, Wilusz J, Wilusz CJ. The highways and byways of mRNA decay. Nat Rev Mol Cell Bio. 2007;8(2):113–26.

2. Weake VM, Workman JL. Inducible gene expression: diverse regulatory mechanisms. Nat Rev Genet. 2010;11(6):426–37.

3. Furlan M, de Pretis S, Pelizzola M. Dynamics of transcriptional and post-transcriptional regulation. Brief Bioinform. 2021;22(4).

4. Rabani M, Levin JZ, Fan L, Adiconis X, Raychowdhury R, Garber M, et al. Metabolic labeling of RNA uncovers principles of RNA production and degradation dynamics in mammalian cells. Nat Biotechnol. 2011;29(5):436–U237.

5. Rabani M, Raychowdhury R, Jovanovic M, Rooney M, Stumpo DJ, Pauli A, et al. High-Resolution Sequencing and Modeling Identifies Distinct Dynamic RNA Regulatory Strategies. Cell. 2014;159(7):1698–710.

6. Dolken L, Ruzsics Z, Radle B, Friedel CC, Zimmer R, Mages J, et al. High-resolution gene expression profiling for simultaneous kinetic parameter analysis of RNA synthesis and decay. RNA. 2008;14(9):1959–72.

7. Windhager L, Bonfert T, Burger K, Ruzsics Z, Krebs S, Kaufmann S, et al. Ultrashort and progressive 4sU-tagging reveals key characteristics of RNA processing at nucleotide resolution. Genome Res. 2012;22(10):2031–42.

8. de Pretis S, Kress T, Morelli MJ, Melloni GE, Riva L, Amati B, et al. INSPEcT: a computational tool to infer mRNA synthesis, processing and degradation dynamics from RNA- and 4sU-seq time course experiments. Bioinformatics. 2015;31(17):2829–35.

9. Duffy EE, Schofield JA, Simon MD. Gaining insight into transcriptome-wide RNA population dynamics through the chemistry of 4-thiouridine. Wires Rna. 2019;10(1).

10. Wada T, Becskei A. Impact of Methods on the Measurement of mRNA Turnover. Int J Mol Sci. 2017;18(12).

11. Schofield JA, Duffy EE, Kiefer L, Sullivan MC, Simon MD. TimeLapse-seq: adding a temporal dimension to RNA sequencing through nucleoside recoding. Nat Methods. 2018;15(3):221–5.

12. Herzog VA, Reichholf B, Neumann T, Rescheneder P, Bhat P, Burkard TR, et al. Thiol-linked alkylation of RNA to assess expression dynamics. Nat Methods. 2017;14(12):1198-+.

13. Riml C, Amort T, Rieder D, Gasser C, Lusser A, Micura R. Osmium-Mediated Transformation of 4-Thiouridine to Cytidine as Key To Study RNA Dynamics by Sequencing. Angew Chem Int Ed Engl. 2017;56(43):13479–83.

14. Jurges C, Dolken L, Erhard F. Dissecting newly transcribed and old RNA using GRAND-SLAM. Bioinformatics. 2018;34(13):i218–i26.

15. Luo Y, Schofield JA, Simon MD, Slavoff SA. Global Profiling of Cellular Substrates of Human Dcp2. Biochemistry. 2020.

16. Uvarovskii A, Dieterich C. pulseR: Versatile computational analysis of RNA turnover from metabolic labeling experiments. Bioinformatics. 2017;33(20):3305–7.

17. Stark R, Grzelak M, Hadfield J. RNA sequencing: the teenage years. Nat Rev Genet. 2019;20(11):631–56.

18. Berge KVd, Hembach KM, Soneson C, Tiberi S, Clement L, Love MI, et al. RNA Sequencing Data: Hitchhiker’s Guide to Expression Analysis. Annual Review of Biomedical Data Science. 2019;2(1):139–73.

19. Costa-Silva J, Domingues D, Lopes FM. RNA-Seq differential expression analysis: An extended review and a software tool. Plos One. 2017;12(12).

20. Li WV, Li JJ. Modeling and analysis of RNA-seq data: a review from a statistical perspective. Quant Biol. 2018;6(3):195–209.

21. Katahira K. How hierarchical models improve point estimates of model parameters at the individual level. J Math Psychol. 2016;73:37–58.

22. Conesa A, Madrigal P, Tarazona S, Gomez-Cabrero D, Cervera A, McPherson A, et al. A survey of best practices for RNA-seq data analysis (vol 17, 13, 2016). Genome Biol. 2016;17.

23. Love MI, Huber W, Anders S. Moderated estimation of fold change and dispersion for RNA-seq data with DESeq2. Genome Biol. 2014;15(12).

24. Robinson MD, McCarthy DJ, Smyth GK. edgeR: a Bioconductor package for differential expression analysis of digital gene expression data. Bioinformatics. 2010;26(1):139–40.

25. Ritchie ME, Phipson B, Wu D, Hu Y, Law CW, Shi W, et al. limma powers differential expression analyses for RNA-sequencing and microarray studies. Nucleic Acids Res. 2015;43(7):e47.

26. Thiecke MJ, Wutz G, Muhar M, Tang W, Bevan S, Malysheva V, et al. Cohesin-Dependent and -Independent Mechanisms Mediate Chromosomal Contacts between Promoters and Enhancers. Cell Rep. 2020;32(3):107929.

27. Biancon G, Joshi P, Zimmer JT, Hunck T, Gao Y, Lessard MD, et al. Precision analysis of mutant U2AF1 activity reveals deployment of stress granules in myeloid malignancies. Mol Cell. 2022;82(6):1107–22 e7.

28. Muhar M, Ebert A, Neumann T, Umkehrer C, Jude J, Wieshofer C, et al. SLAM-seq defines direct gene-regulatory functions of the BRD4-MYC axis. Science. 2018;360(6390):800–5.

29. Luo Y, Schofield JA, Na Z, Hann T, Simon MD, Slavoff SA. Discovery of cellular substrates of human RNA-decapping enzyme DCP2 using a stapled bicyclic peptide inhibitor. Cell Chem Biol. 2021;28(4):463–74 e7.

30. Na Z, Luo Y, Schofield JA, Smelyansky S, Khitun A, Muthukumar S, et al. The NBDY Microprotein Regulates Cellular RNA Decapping. Biochemistry. 2020;59(42):4131–42.

31. Olivero CE, Martinez-Terroba E, Zimmer J, Liao C, Tesfaye E, Hooshdaran N, et al. p53 Activates the Long Noncoding RNA Pvt1b to Inhibit Myc and Suppress Tumorigenesis. Mol Cell. 2020;77(4):761–74 e8.

32. Zimmer JT, Rosa-Mercado NA, Canzio D, Steitz JA, Simon MD. STL-seq reveals pause-release and termination kinetics for promoter-proximal paused RNA polymerase II transcripts. Mol Cell. 2021;81(21):4398–412 e7.

33. Gasser C, Delazer I, Neuner E, Pascher K, Brillet K, Klotz S, et al. Thioguanosine Conversion Enables mRNA-Lifetime Evaluation by RNA Sequencing Using Double Metabolic Labeling (TUC-seq DUAL). Angew Chem Int Edit. 2020;59(17):6881–6.

34. Kiefer L, Schofield JA, Simon MD. Expanding the Nucleoside Recoding Toolkit: Revealing RNA Population Dynamics with 6-Thioguanosine. J Am Chem Soc. 2018;140(44):14567–70.

35. Reichholf B, Herzog VA, Fasching N, Manzenreither RA, Sowemimo I, Ameres SL. Time-Resolved Small RNA Sequencing Unravels the Molecular Principles of MicroRNA Homeostasis. Mol Cell. 2019;75(4):756–68 e7.

36. Lonnstedt I, Speed T. Replicated microarray data. Stat Sinica. 2002;12(1):31–46.

37. Wu H, Wang C, Wu Z. A new shrinkage estimator for dispersion improves differential expression detection in RNA-seq data. Biostatistics. 2013;14(2):232–43.

38. Gelman A, Lee D, Guo JQ. Stan: A Probabilistic Programming Language for Bayesian Inference and Optimization. J Educ Behav Stat. 2015;40(5):530–43.

39. Gelman A. Bayesian data analysis. Third edition. ed. Boca Raton: CRC Press; 2014. xiv, 661 pages p.

40. Brooks S, Gelman A, Jones GL, Meng XL. Handbook of Markov Chain Monte Carlo Preface. Ch Crc Handb Mod Sta. 2011:Xix–Xx.

41. Matthews BW. Comparison of the predicted and observed secondary structure of T4 phage lysozyme. Biochim Biophys Acta. 1975;405(2):442–51.

42. Chicco D, Jurman G. The advantages of the Matthews correlation coefficient (MCC) over F1 score and accuracy in binary classification evaluation. Bmc Genomics. 2020;21(1).

43. Dunckley T, Parker R. The DCP2 protein is required for mRNA decapping in Saccharomyces cerevisiae and contains a functional MutT motif. Embo J. 1999;18(19):5411–22.

44. Chen KF, Hu Z, Xia Z, Zhao DY, Li W, Tyler JK. The Overlooked Fact: Fundamental Need for Spike-In Control for Virtually All Genome-Wide Analyses. Mol Cell Biol. 2016;36(5):662–7.

45. Sun M, Schwalb B, Schulz D, Pirkl N, Etzold S, Lariviere L, et al. Comparative dynamic transcriptome analysis (cDTA) reveals mutual feedback between mRNA synthesis and degradation. Genome Res. 2012;22(7):1350–9.

46. Haimovich G, Medina DA, Causse SZ, Garber M, Millan-Zambrano G, Barkai O, et al. Gene Expression Is Circular: Factors for mRNA Degradation Also Foster mRNA Synthesis. Cell. 2013;153(5):1000–11.

47. Abernathy E, Gilbertson S, Alla R, Glaunsinger B. Viral Nucleases Induce an mRNA Degradation-Transcription Feedback Loop in Mammalian Cells. Cell Host Microbe. 2015;18(2):243–53.

48. Gilbertson S, Federspiel JD, Hartenian E, Cristea IM, Glaunsinger B. Changes in mRNA abundance drive shuttling of RNA binding proteins, linking cytoplasmic RNA degradation to transcription. Elife. 2018;7.

49. Berry S, Muller M, Rai A, Pelkmans L. Feedback from nuclear RNA on transcription promotes robust RNA concentration homeostasis in human cells. Cell Syst. 2022.

50. Faucillion ML, Johansson AM, Larsson J. Modulation of RNA stability regulates gene expression in two opposite ways: through buffering of RNA levels upon global perturbations and by supporting adapted differential expression. Nucleic Acids Res. 2022;50(8):4372–88.

51. Luo Y, Na ZK, Slavoff SA. P-Bodies: Composition, Properties, and Functions. Biochemistry. 2018;57(17):2424–31.

52. Hubstenberger A, Courel M, Benard M, Souquere S, Ernoult-Lange M, Chouaib R, et al. P-Body Purification Reveals the Condensation of Repressed mRNA Regulons. Mol Cell. 2017;68(1):144–57 e5.

53. Fu Y, Dominissini D, Rechavi G, He C. Gene expression regulation mediated through reversible m(6)A RNA methylation. Nat Rev Genet. 2014;15(5):293–306.

54. Batista PJ, Molinie B, Wang JK, Qu K, Zhang JJ, Li LJ, et al. m(6)A RNA Modification Controls Cell Fate Transition in Mammalian Embryonic Stem Cells. Cell Stem Cell. 2014;15(6):707–19.

55. Furlan M, Galeota E, Del Gaudio N, Dassi E, Caselle M, de Pretis S, et al. Genome-wide dynamics of RNA synthesis, processing, and degradation without RNA metabolic labeling. Genome Res. 2020;30(10):1492–507.

56. McElreath R. Statistical rethinking: a Bayesian course with examples in R and Stan. 2. ed. Boca Raton: Taylor and Francis, CRC Press; 2020. pages cm p.

57. Betancourt MJ, Girolami M. Hamiltonian Monte Carlo for Hierarchical Models [Internet]. arxiv. 2013;Available from: https://arxiv.org/abs/1312.0906.

58. Redner RA, Walker HF. Mixture Densities, Maximum-Likelihood and the Em Algorithm. Siam Rev. 1984;26(2):195–237.

59. Smyth GK. Linear models and empirical bayes methods for assessing differential expression in microarray experiments. Stat Appl Genet Mol Biol. 2004;3:Article3.

60. Benjamini Y, Hochberg Y. Controlling the False Discovery Rate - a Practical and Powerful Approach to Multiple Testing. J R Stat Soc B. 1995;57(1):289–300.

61. Stephens M. False discovery rates: a new deal. Biostatistics. 2017;18(2):275–94.

62. Xu H, Luo X, Qian J, Pang X, Song J, Qian G, et al. FastUniq: a fast de novo duplicates removal tool for paired short reads. PLoS One. 2012;7(12):e52249.

63. Martin M. Cutadapt removes adapter sequences from high-throughput sequencing reads. EMBnetjournal. 2011;12(4):357–60.

64. Zhang Y, Park C, Bennett C, Thornton M, Kim D. Rapid and accurate alignment of nucleotide conversion sequencing reads with HISAT-3N. Genome Res. 2021.

65. Anders S, Pyl PT, Huber W. HTSeq--a Python framework to work with high-throughput sequencing data. Bioinformatics. 2015;31(2):166–9.

66. Li H, Handsaker B, Wysoker A, Fennell T, Ruan J, Homer N, et al. The Sequence Alignment/Map format and SAMtools. Bioinformatics. 2009;25(16):2078–9.

67. Narasimhan V, Danecek P, Scally A, Xue Y, Tyler-Smith C, Durbin R. BCFtools/RoH: a hidden Markov model approach for detecting autozygosity from next-generation sequencing data. Bioinformatics. 2016;32(11):1749–51.

68. Tange O. GNU Parallel 20182018. Available from: https://doi.org/10.5281/zenodo.1146014.

69. Korotkevich G. Fast gene set enrichment analysis [Internet]. biorxiv. 2021;Available from: http://biorxiv.org/content/early/2016/06/20/060012.

