## Additional file 1: Supplemental Figures for "Differential kinetic analysis using nucleotide recoding RNA-seq and bakR"

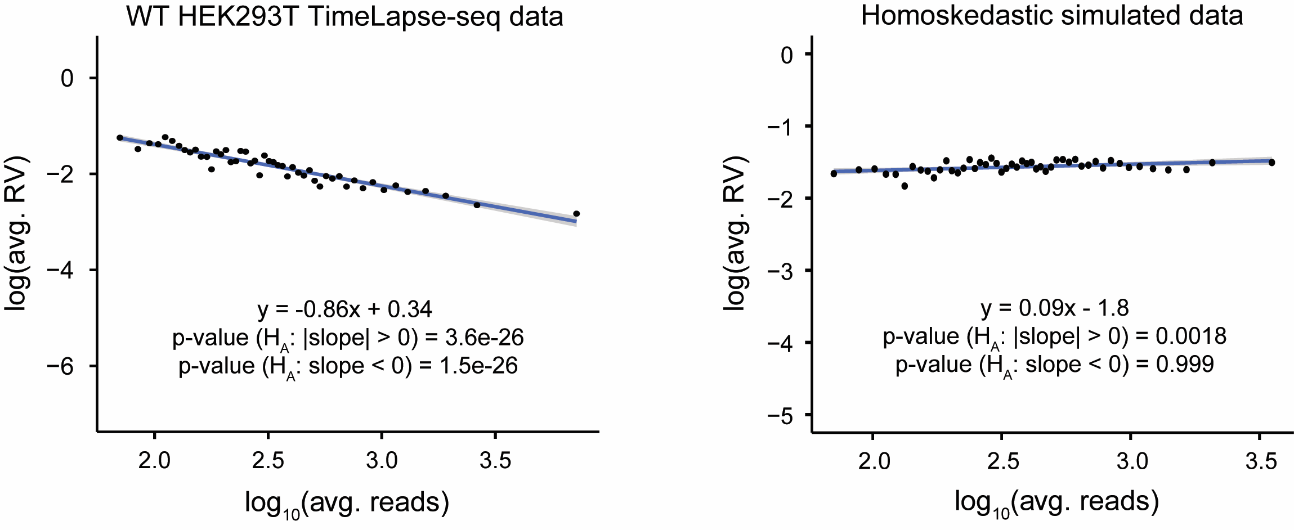

**Fig. S1: Analysis of depth vs. RV trend is unbiased.** **Left:** Replication of data shown in Fig. 3C. **Right:** Identical analysis as performed in Fig. 3C but on simulated homoskedastic data where there is no relationship between RV and sequencing depth.

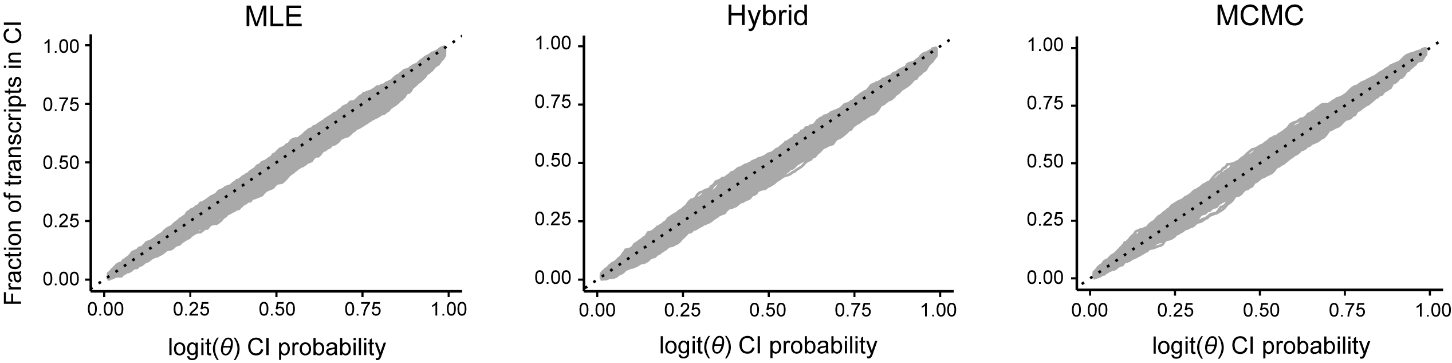

**Fig. S2: logit(*θ*) calibration analysis for the three implementations of the novel hierarchical model.** All implementations are well calibrated, with the Hybrid model showing slight underestimation of *θ* uncertainty. Data is 500 lines obtained from bootstrapped samples of 1000 transcripts from a total of 10000 transcripts.

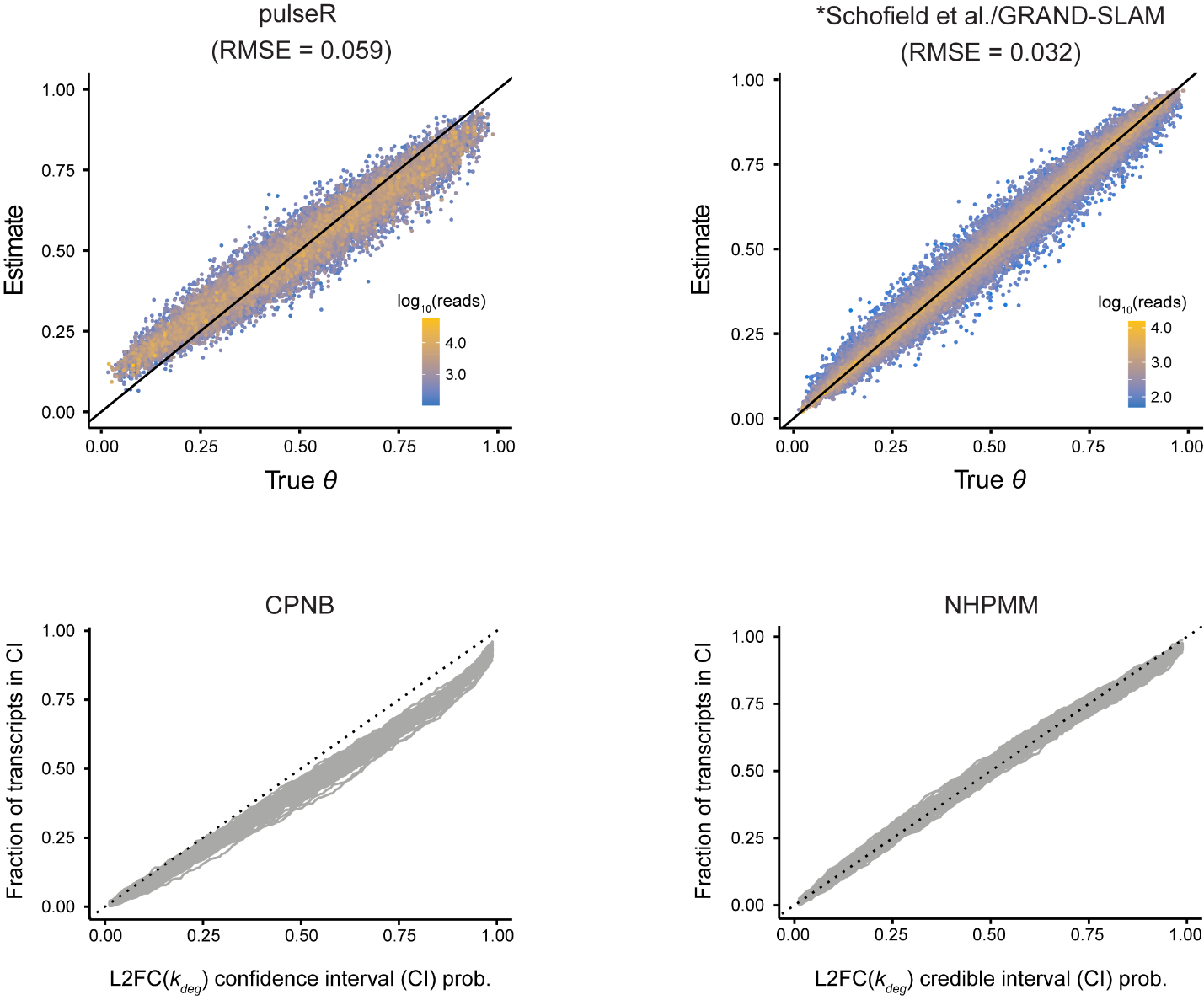

**Fig. S3: Simulated data validation of extensions of existing models of NR-seq data.** **Top:** Accuracy of $\theta$ estimates for existing models. Diagonal line is y = x. *Schofield et al./GRAND-SLAM model is a U-content adjusted Poisson rather than the original binomial mixture model. See **Methods** and Additional file 2: Supplemental Methods for details. Note, the pulseR RMSEs are not directly comparable with those for the other models/implementations, as pulseR only provides replicate average $\theta$ estimates, whereas all other models provide estimates of $\theta$ in each replicate and is what is compared to simulated truth here and Fig. 4B. **Bottom:** L2FC($k_{\deg})$calibration for both implementations of the existing models. The x-axis represents the coverage probability of the L2FC($k_{\deg})$ credible/confidence interval and the y-axis is the percentage of transcripts who’s true L2FC($k_{\deg})$ fall within the credible/confidence interval. 500 lines obtained from bootstrapped samples of 1000 transcripts from a total of 10000 transcripts are depicted for each implementation.

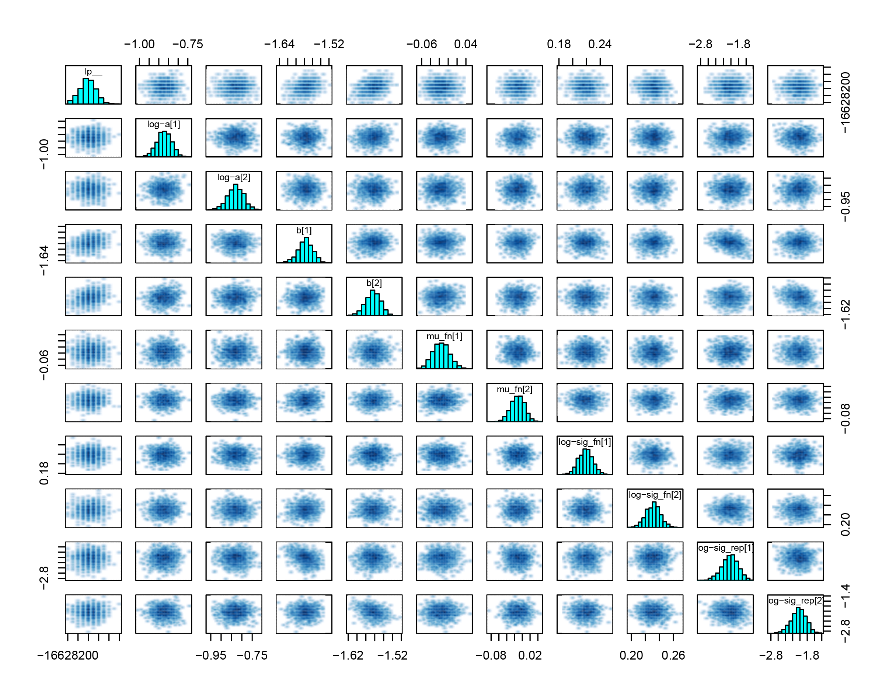

**Fig. S4: Pairs plots for hierarchical hyperparameters indicate unbiased estimation.** Pairs plots of marginal posterior distributions are shown for the hyper-parameters labeled along the diagonal. The lack of correlation between any of the marginals indicates identifiability of all parameters.

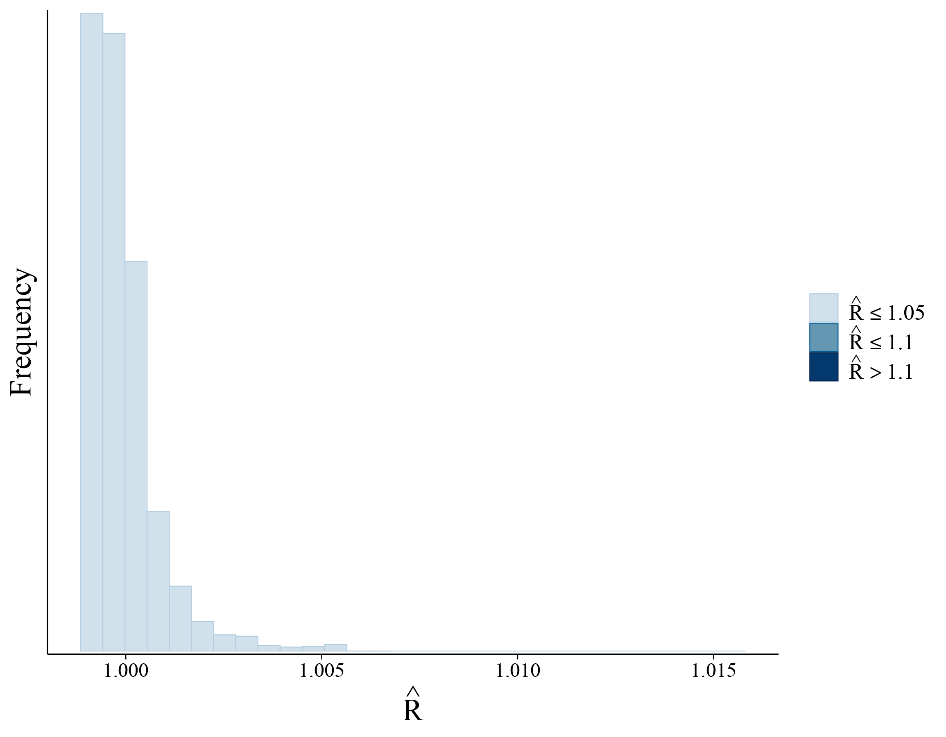

**Fig. S5:** $\hat{\mathbf{R}}$ **distribution indicates model convergence.** The $\hat{R}$ is a quantification of the extent to which multiple independent Markov chains converged to the same posterior samples for a parameter. An $\hat{R}$ closer to 1 is better.

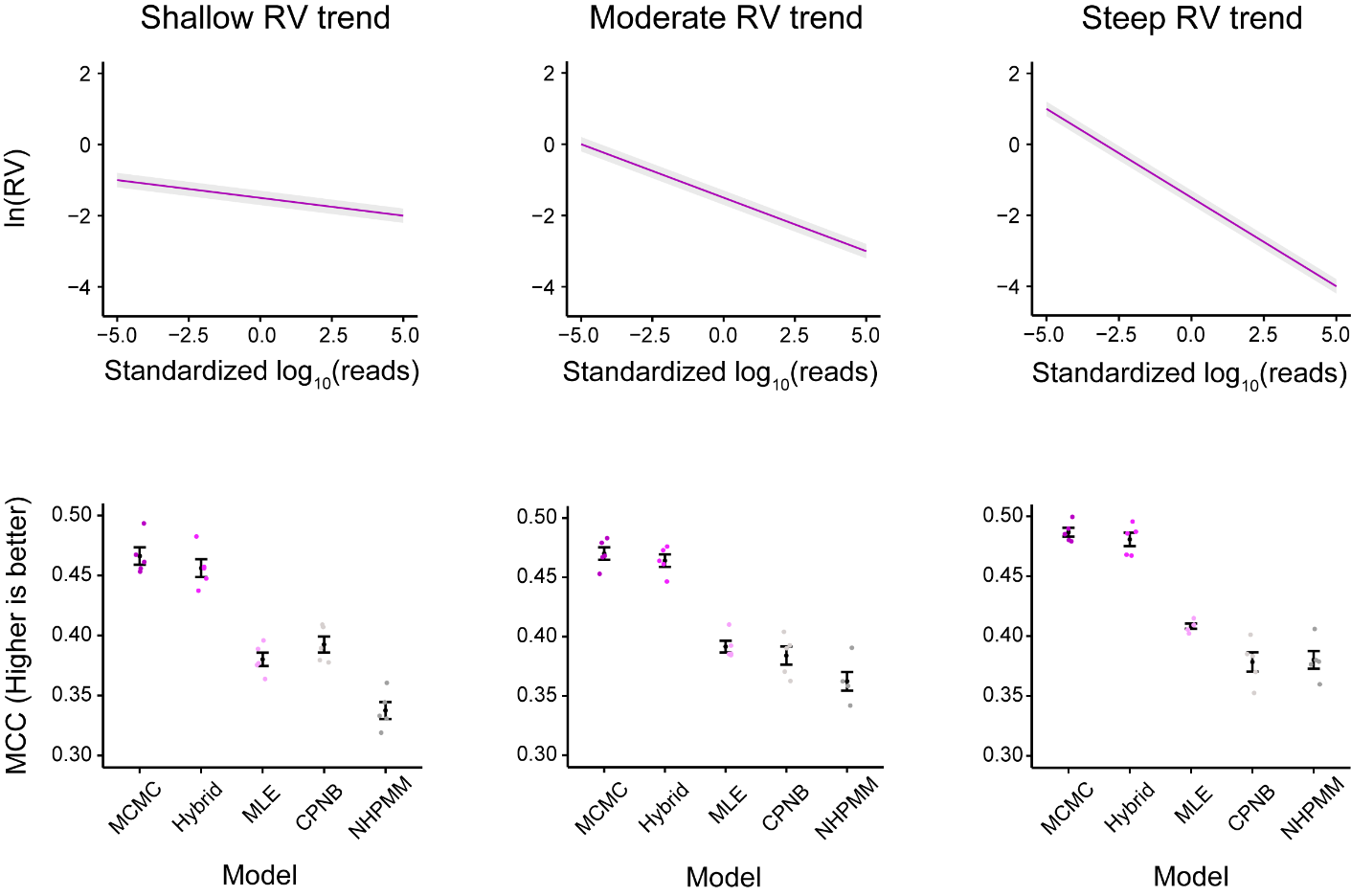

**Fig. S6: MCCs confirm performance hierarchy.** **Top:** Simulated ln(RV) vs. standardized read count trend. ln(RV) for each transcript is drawn from a normal distribution centered on the trend with standard deviation of 0.05. Gray bar around the trend line represents the range for 99% of simulated replicate variabilities. **Bottom:** Matthew’s correlation coefficient (MCC) plotted for each simulation replicate and a p_adj_ significance cutoff of 0.05. x-axis jitter applied to all points to limit overlaps. Black points are averaged MCC values and error bars represent ± one standard error.

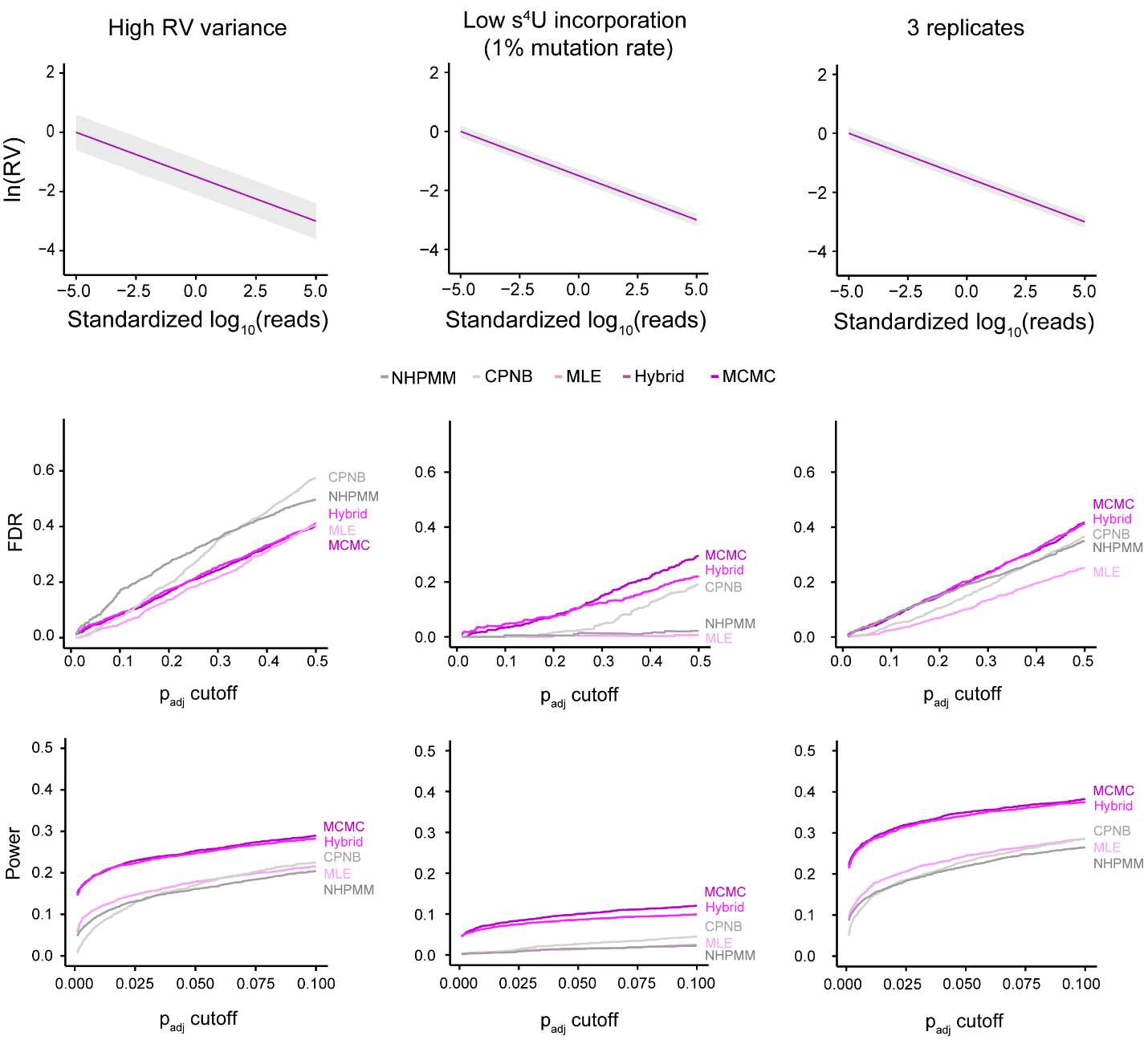

**Fig. S7: Additional simulations show that bakR implementations outperform extensions of existing models.** **Top:** Simulated ln(RV) vs. standardized read count trend. The gray bar around the trend lines represent the range for 99% of simulated replicate variabilities. The left and center columns are results for simulations of two biological replicates; right column is for simulation of three biological replicates. The center column is a simulation of low s^4^U incorporation (1%, as opposed to the 5% incorporation rate simulated in Fig. 5 data). **Middle:** False discovery rate (FDR) as a function of significance cutoff for all implementations. **Bottom:** Power as a function of significance cutoff for all implementations. The average FDR and Power for five simulation replicates are plotted at each cutoff

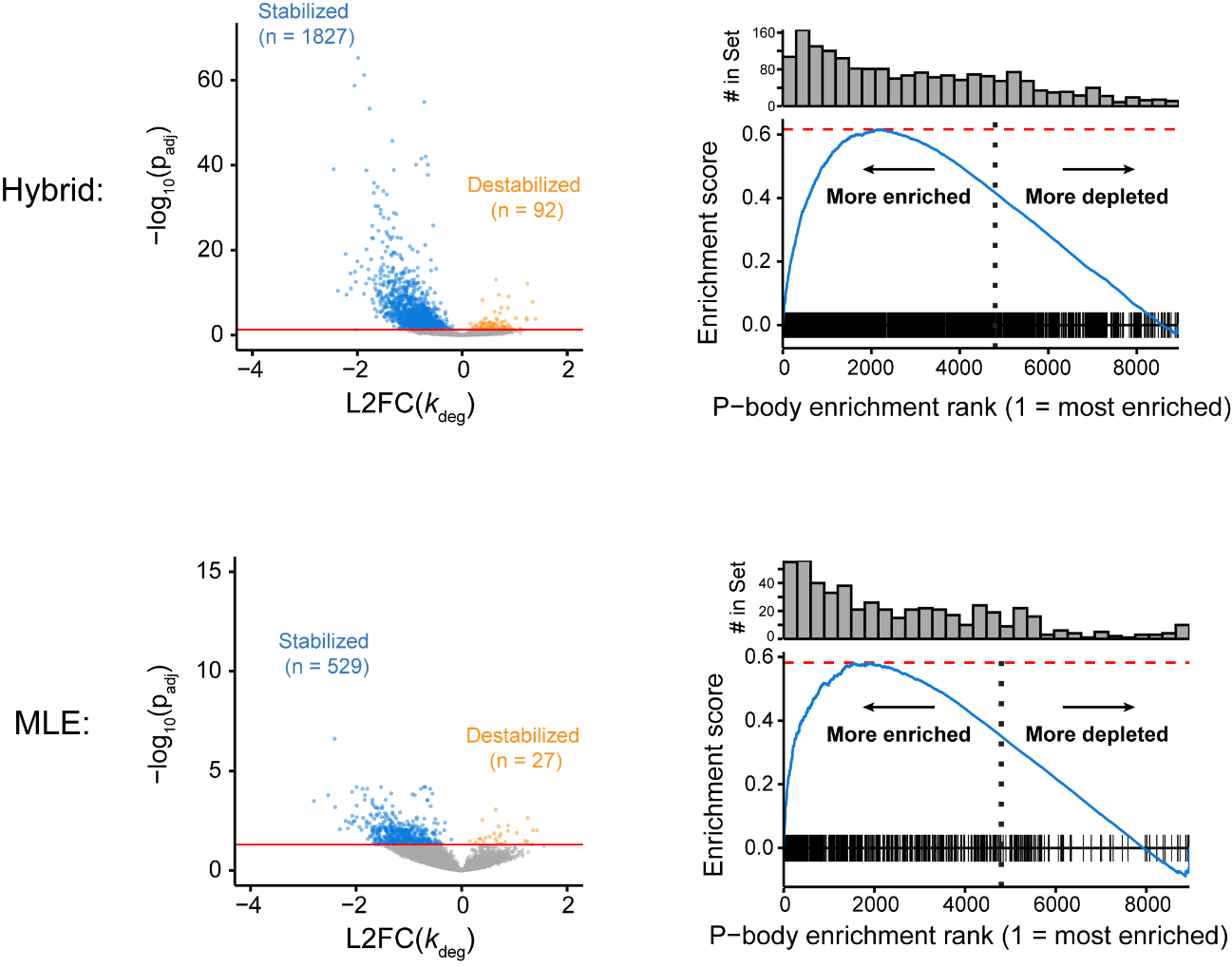

**Fig. S8: All bakR implementations reveal true biological signal in TimeLapse-seq data.** **Left:** Volcano plots from the comparison of *DCP2* KO and WT HEK293T cells using data from Luo et al. The top volcano plot is from analysis with the Hybrid implementation and the bottom volcano plot is from analysis with the MLE implementation. **Right:** GSEA analysis using the transcripts identified as stabilized as the gene set of interest and P-body enrichment scores as the scoring function.

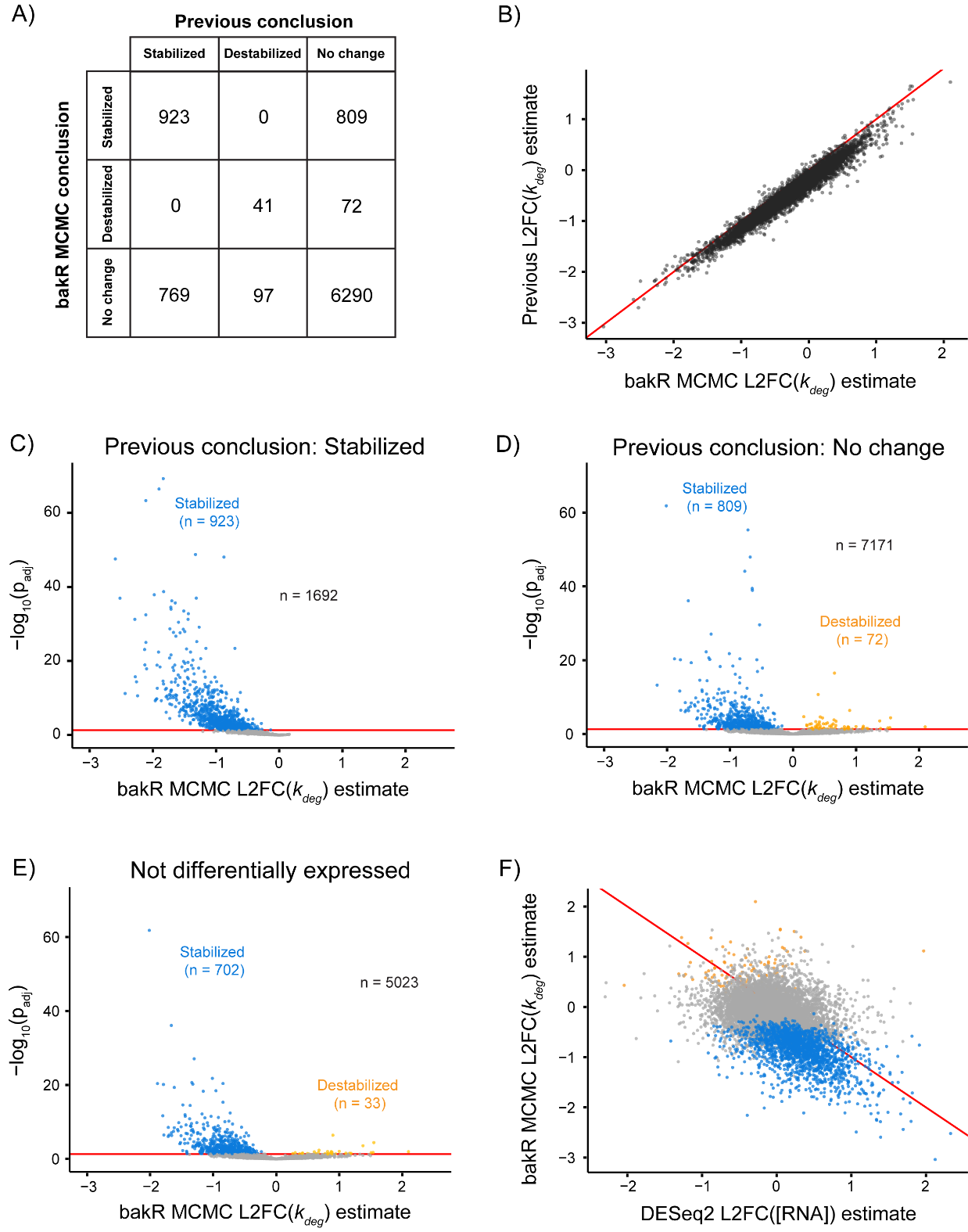

**Fig. S9: bakR provides an improved list of differentially stable transcripts independent of differential expression.** **A)** Comparison of conclusions using the previous anlaysis strategy of Luo et al. and bakR’s MCMC implementation. **B)** Correlation between estimates of the log-2 fold change in the degradation rate constant (L2FC(*k*_deg_); comparing *DCP2* KO to WT) from the previous analysis strategy and bakR’s MCMC implementation. Red line is y = x. **C)** bakR MCMC volcano plot for transcripts identified as stabilized with the previous analysis strategy. **D)** bakR MCMC volcano plot for the transcripts identified as not stabilized/destabilized with the previous analysis strategy. **E)** bakR MCMC volcano plot for the transcripts identified as not differentially expressed by the previous analysis strategy’s use of DESeq2. **F)** Correlation between differential expression and differential stability. Red line is y = x.

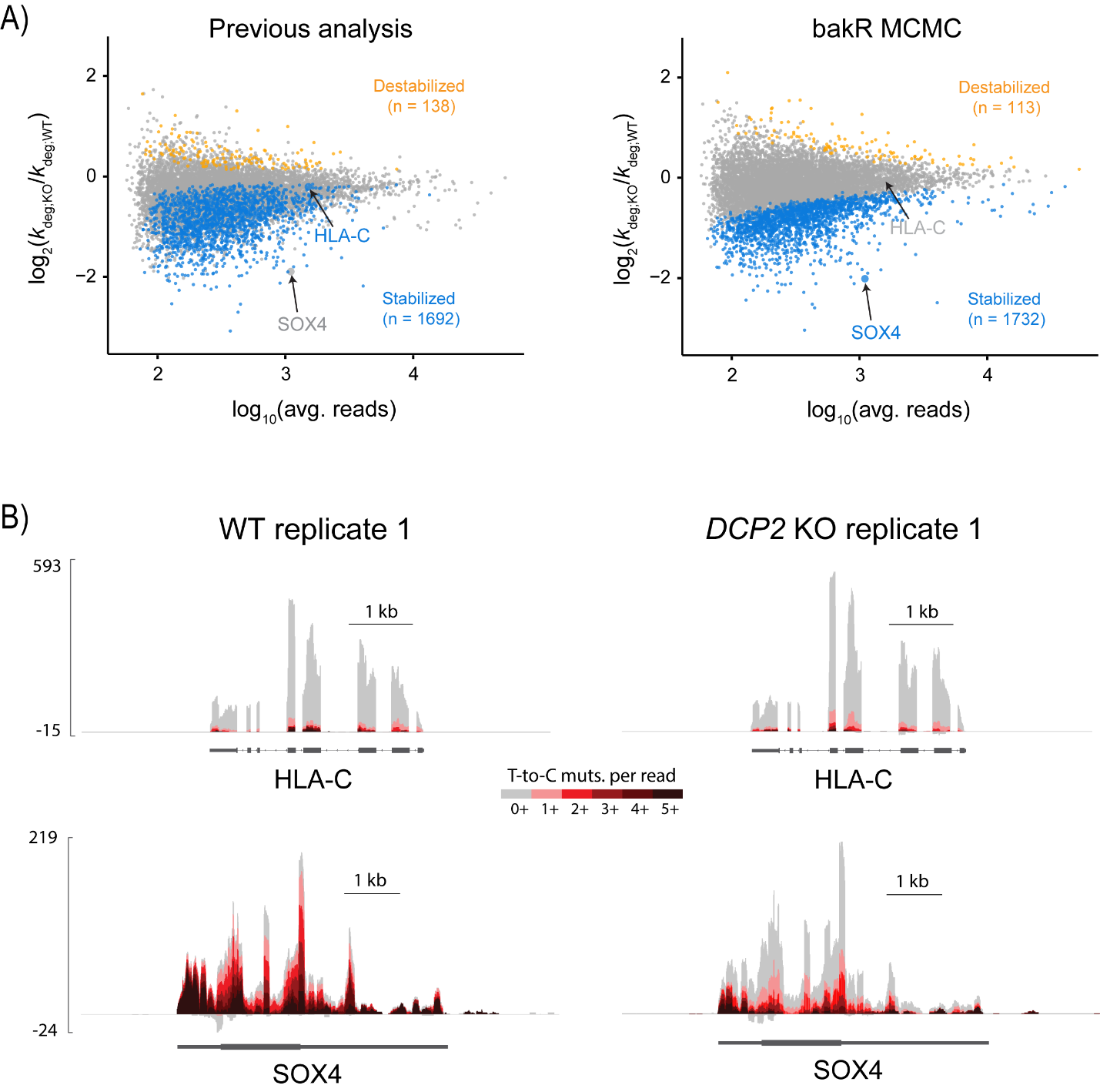

**Fig. S10: bakR provides an improved list of *DCP2* KO stabilized transcripts.** **A)** Same data as in Fig. 6C but with two transcripts highlighted. HLA-C is an example of a low confidence, trivial effect size transcript that the previous anlaysis identified as stabilized and bakR’s MCMC implementation identified as not significant. SOX4 is an example of a transcript that the previous anlaysis did not consider because it was not identified as differentially expressed by DEseq2, despite having strong evidence for stabilization. **B)** Sequencing tracks for the two representative transcripts from one replicate of the two conditions (WT and *DCP2* KO). Tracks are colored by the mutational content of sequencing reads

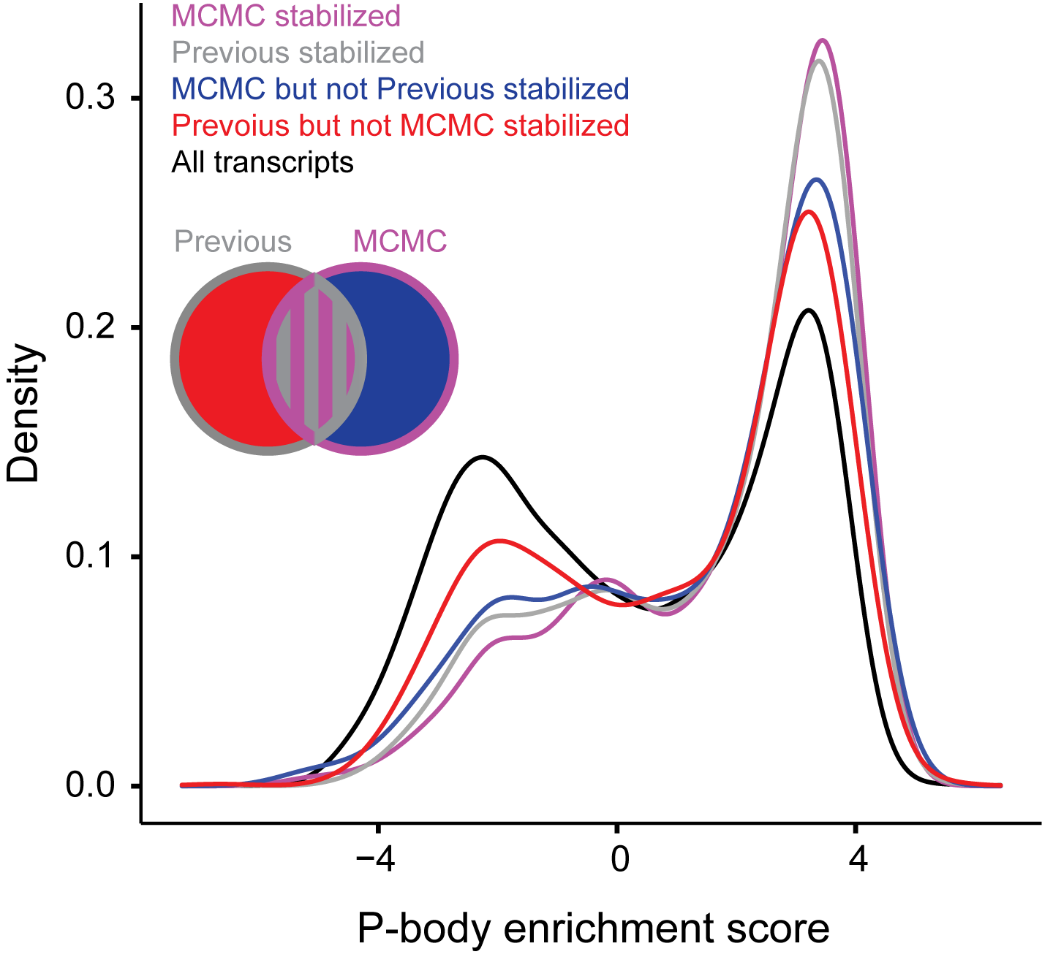

**Fig. S11: The bakR list of DCP2 KO stabilized transcripts demonstrate an increased signal for P-body enrichment compared with the previous anlaysis.** The x-axis is derived from the scoring function used for GSEA. Positive scores represent transcripts that are enriched in P-bodies. Negative scores represent transcripts that are depleted from P-bodies. MCMC stabilized is the set of transcripts identified as stabilized (FDR = 0.05) by bakR’s MCMC implementation. Previous stabilized is the set of transcripts identified as stabilized by the previous analysis strategy. MCMC but not Previous stabilized are those transcripts in the MCMC stabilized set and not the Previous stabilized set. Previous but not MCMC stabilized is those transcripts in the Previous stabilized set and not the MCMC stabilized set. The Venn diagram depicts the relationships between these four sets.

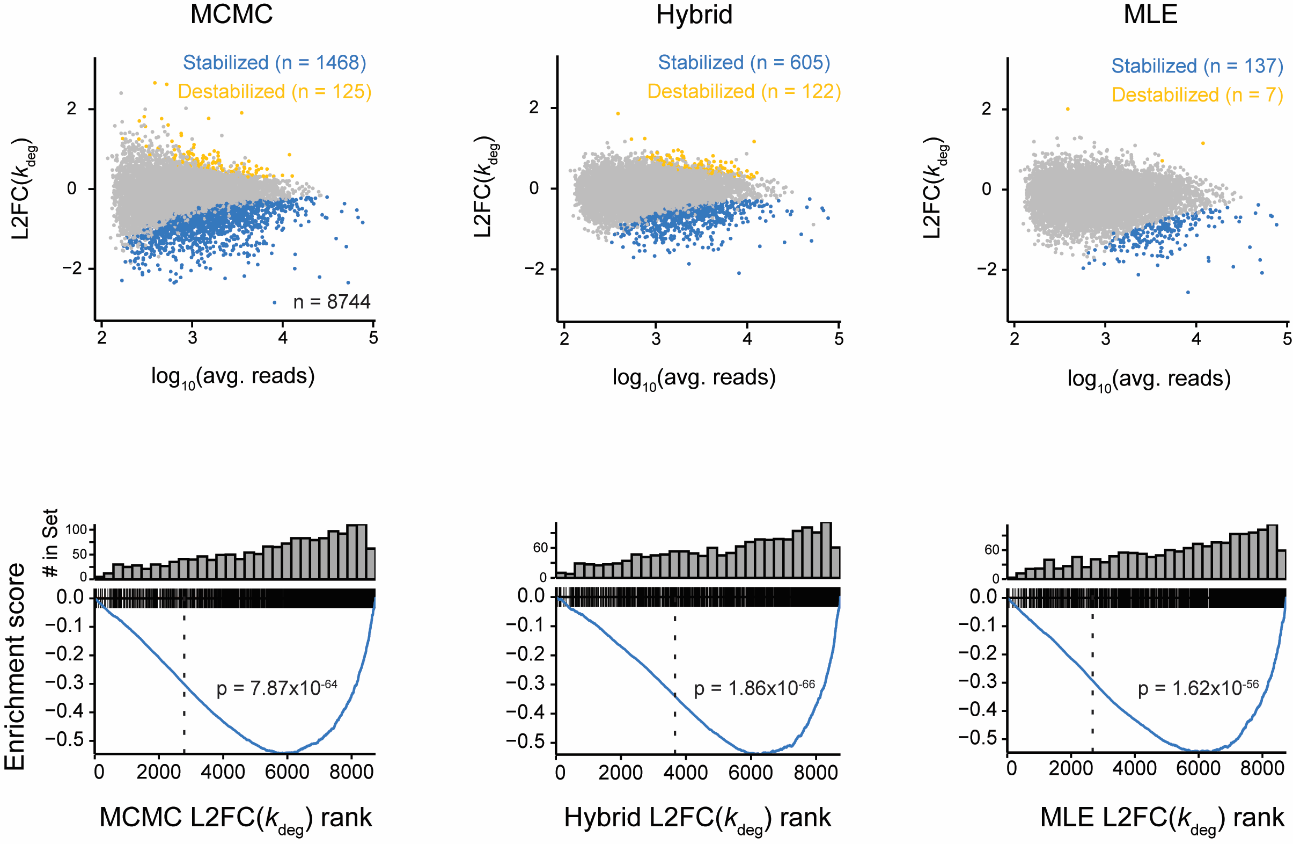

**Fig. S12: bakR identifies biological signal in SLAM-seq datasets. Top**: MA plots for comparison of Mettl3 KO and WT mESC SLAM-seq data. Results for all three bakR implementations shown. **Bottom:** GSEA using transcripts previously identified as being targets of Mettl3 as the gene set of interest and L2FC(*k_deg_*) z-scores as the scoring function.

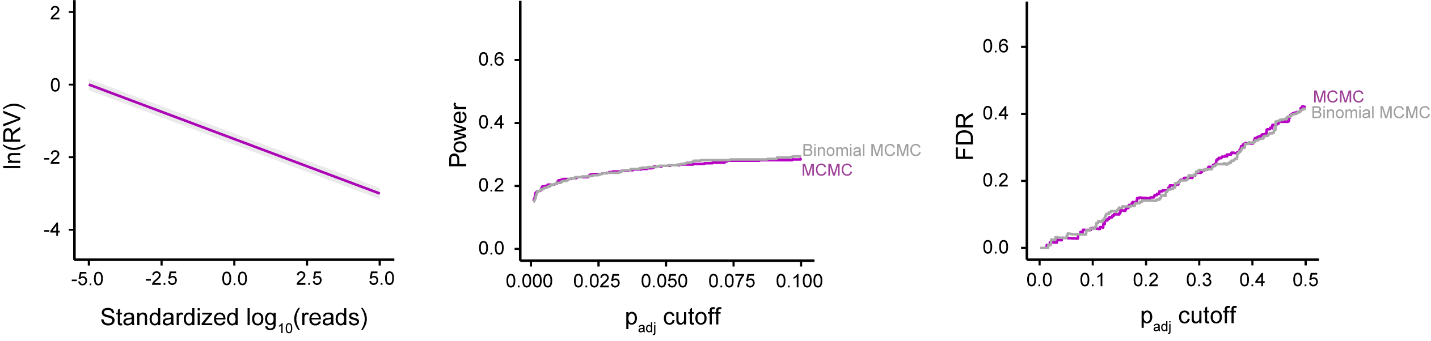

**Fig. S13: Binomial and U-content adjusted Poisson mixture models perform equally well. Left:** Replicate variability vs. read count trend simulated. **Center:** Statistical power as a function of significance cutoff for both models. Binomial MCMC is identical to bakR’s MCMC implementation except that mutations are modeled as coming from a binomial distribution rather than a U-content adjusted Poisson distribution. **Right:** FDR as a function of significance cutoff.

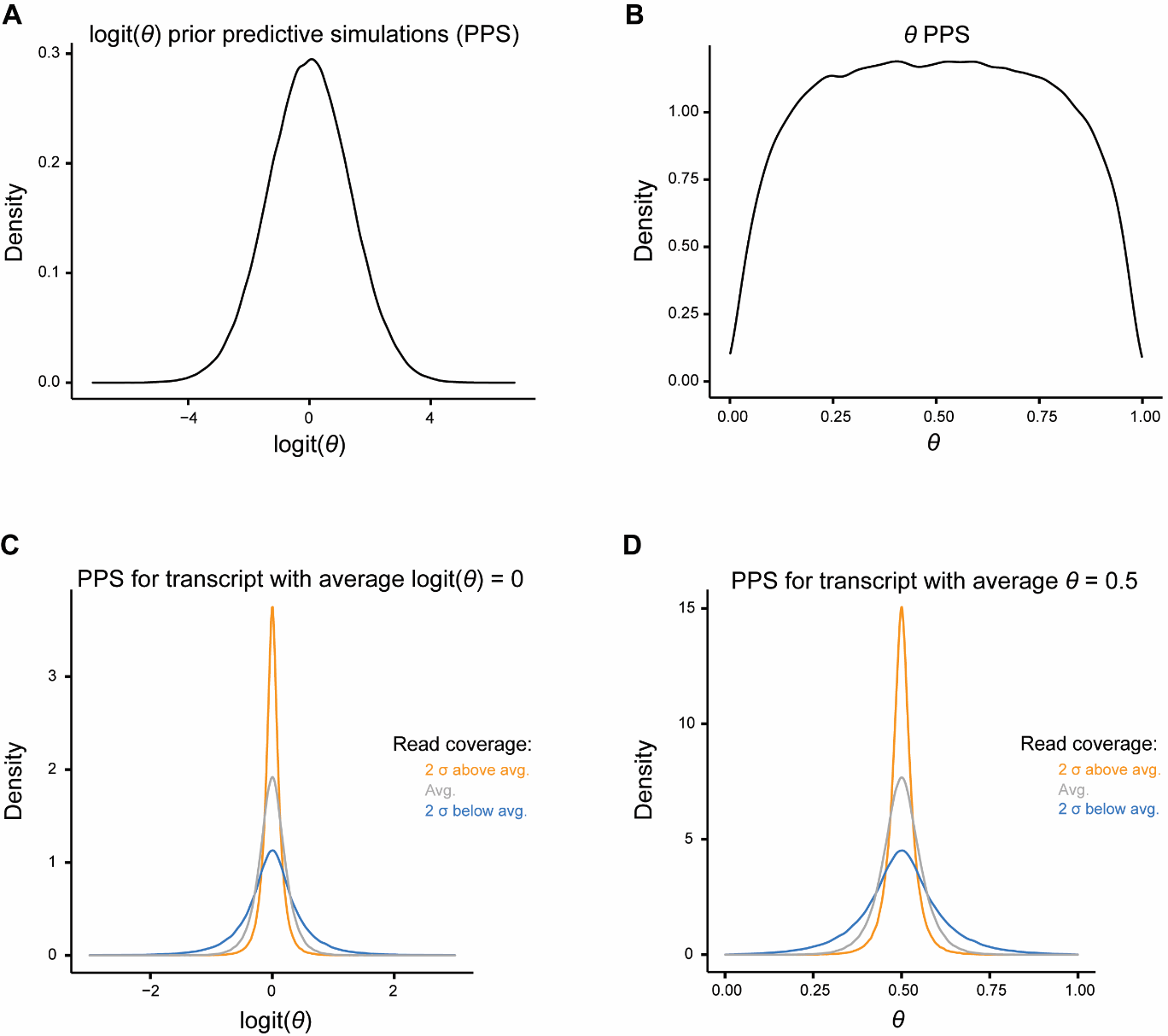

**Fig. S14: Fraction new prior predictive distributions used in bakR represent realistic values. A)** Prior distribution of replicate logit(*θ*) estimates. The simulation is effectively of three biological replicates for 100,000 transcripts and represents the model’s per-transcript per-replicate logit(*θ*) prior. **B)** Same simulation as **A)** but on the natural rather than logit scale. **C)** Prior predictive distribution of replicate logit(*θ*) for a transcript with an average logit(*θ*) of 0 (average *θ* of 0.5). Simulation shown for theoretical low, average, and high coverage transcript. **D)** Same as **C)** but on the natural rather than logit scale

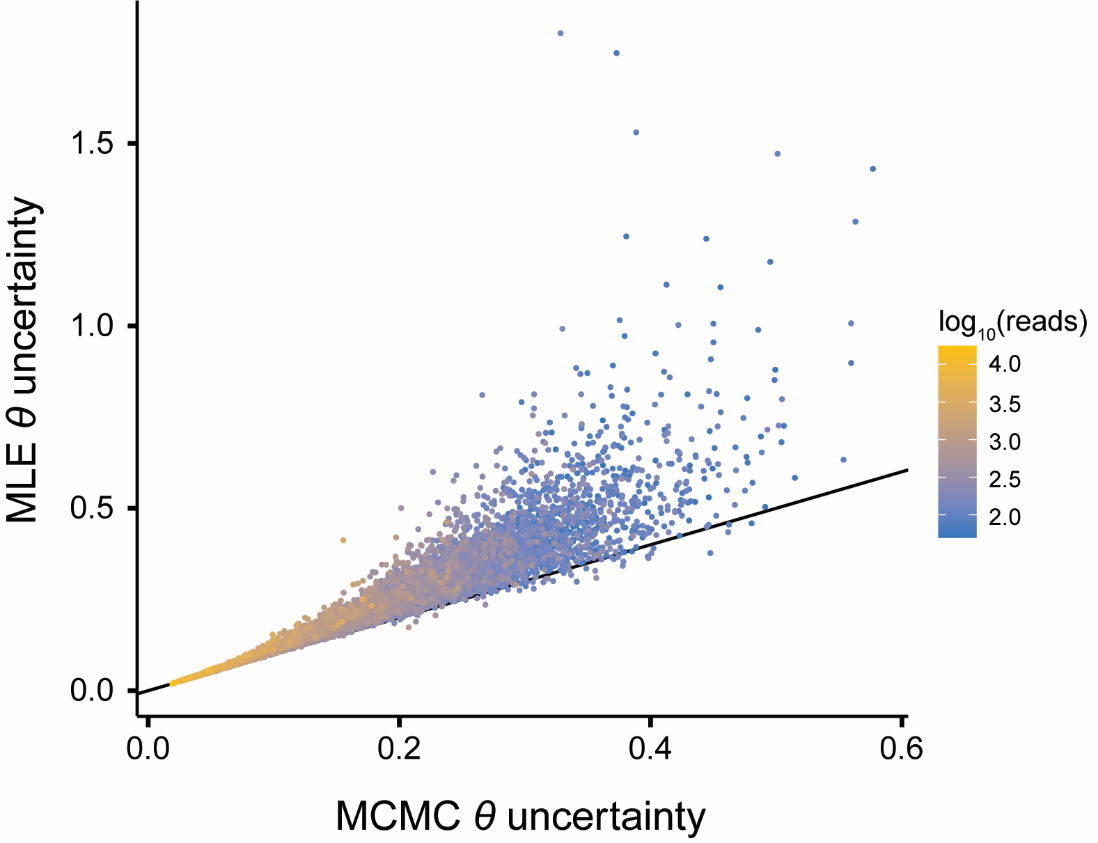

**Fig. S15: MLE *θ* uncertainty is well quantified at high sequencing depth and is conservative at low seqeuncing depths.** Comparison of Fisher Information *θ* uncertainty estimate used by the MLE implementation to the *θ* posterior standard deviation estimated by the MCMC implementation.

**Supplemental Table:**

|  | **2 replicate simulation;**  **moderate heteroskedasticity** | **2 replicate Luo et al. TimeLapse-seq dataset** | **3 replicate simulation** |
| --- | --- | --- | --- |
| **MLE** | 4m 14s | 5m 15s | 6m 23s |
| **CPNB** | 33 m 8s | 53m 30s | 43 m 24 s |
| **Hybrid** | 1h 56m | 1h 30m | 4 h 13m |
| **NHPMM** | ~2d | ~2d | ~3d |
| **MCMC** | ~2d | ~2d | ~3d |

**Table S1: Representative runtimes of the five implementations tested.** Runtimes shown are for one run of each of the three datasets listed. Analyses were performed on the Ruddle HPC cluster at Yale University using a one CPU core for the MLE and CPNB implementations and two cores for the MCMC, NHPMM, and Hybrid implementations. For the MCMC, NHPMM, and Hybrid implementations, two Markov chains were run in parallel for 3000 iterations (1500 warmup samples). The MLE implementation was run with StanRateEst set to true; all other settings were the default. In general, runtimes for the MLE implementation on the datasets analyzed in this paper were on the order of minutes, for the CPNB implementation they were on the order of tens of minutes to an hour, for the Hybrid implementation they were on the order of hours, and for the NHPMM and MCMC implementations they were on the order of 1-3 days. As the CPNB implementation is the only implementation to not model sequencing read mutational data directly, its runtimes scale more slowly with the number of replicates and transcripts, leading to roughly similar runtimes for all three datasets discussed in this table.
