## Additional file 2: Supplemental Methods for "Differential kinetic analysis using nucleotide recoding RNA-seq and bakR"

**MLE implementation details**

The MLE implementation makes use of the generative model presented in **Methods.** Rather than using Stan to sample from the posterior using Hamiltonian Monte Carlo like in the MCMC implementation, the MLE implementation makes several approximations to increase efficiency. The MLE implementation can be broken down into four parts, each of which will be described in detail, those being mutation rate estimation, $\theta$ estimation and uncertainty quantification in each s^4^U treated sample for each transcript, linear modeling of the replicate variability trend, and approximate partial pooling.

*Mutation rate estimation*

The MLE implementation uses the method of maximum likelihood to estimate $\theta$ (or more specifically logit($\theta$))for each transcript in each sample. Performing maximum likelihood estimation and using the Fisher Information to approximate estimate uncertainty is fundamentally impossible with a mixture model though. To get around the problem of mixture models being singular, the MLE implementation starts by estimating the mutation rates (${\lambda new}_{e}$ and *λold*), using these fixed estimates when finding the $\theta$ MLE for each transcript in each sample, and adding a prior penalization to avoid boundary estimates (i.e., fraction new estimates of 0 or 1) (1). In bakR, there are two options for how to estimate the mutation rates. The default option assumes that some transcripts have a $\theta$ close to 1, and that some have a $\theta$ close to 0. Thus, it uses the highest and lowest average empirical mutation rates (number of mutations in sequencing reads divided by number of Us in the sequencing reads) of transcripts with sufficient sequencing depth (100 reads is the default cutoff, but this cutoff is a parameter which can be changed by the user) as the estimates of ${\lambda new}_{s}$ and *λold* respectively. The lack of s subscript in *λold* is because a single background mutation rate is estimated for all samples. One reason for this is because the background mutation rate is mostly determined by the library prep kit and sequencer that is used, which is typically constant across samples. In addition, if the user has included samples in which no s^4^U feed was conducted (so called -s^4^U controls), then the average empirical mutation rates in the -s^4^U controls can be used to estimate *λold* more accurately. For ${\lambda new}_{s}$ and *λold* estimation, the default setting in bakR is to use the 7 highest and lowest empirical mutation rates, though the user can also change this setting.

The default strategy will provide decent mutation rate estimates if -s^4^U controls are present and if the s^4^U feed time is long enough to ensure that there are a handful of transcripts which are nearly completely labeled. If the feed time is short or no such controls exist, then a second less efficient mutation rate estimation strategy can be used.

The second strategy involves running a small subset of transcripts through the non-hierarchical Poisson mixture model (NHPMM) implemented in Stan to estimate the mutation rate in each sample. The default number of transcripts to use is 25. Transcripts with sequencing depths around the median are randomly chosen, as these transcripts likely have a fraction new around 0.5. This ensures that both new and old reads are present to allow the model to properly fit the two-component mixture model. All analyses of real and simulated data with the MLE implementation presented here used this mutation rate estimation strategy

$logit(\theta)$ *estimation and uncertainty quantification*

With the mutation rates estimated, estimating $\mathrm{logit}(\theta)$ via the method of maximum likelihood becomes a simple 1-dimensional optimization problem. bakR maximizes a prior penalized likelihood using the optim function and the L-BFGS-B method, implemented in the R package stats (2). A prior penalization of a normal distribution with default standard deviation of 1.25 is added to the likelihood function to approximate the regularization provided by partial pooling in the MCMC model. It also largely eliminates estimates at the upper and lower bounds of the provided estimation range, which arise from transcripts with limited or extreme data (e.g., all new or old reads).

To assign a standard error to each $\mathrm{logit}(\theta)$ estimate, we use the asymptotic result that under smoothness conditions on the likelihood function, the sampling distribution of the MLE tends to a normal distribution with standard deviation (3):

$$\sigma_{\mathrm{MLE}} = \frac{1}{\sqrt{n*I(\theta_{\mathrm{MLE}})}}$$

$$I(\theta_{\mathrm{MLE}}) = {E\left[ \frac{d}{d\theta}\log f(X|\theta_{\mathrm{MLE}}) \right]}^{2}; Fisher Information$$

where $\log f(X|\theta_{\mathrm{MLE}})$ is the 1-dimensional log-likelihood function evaluated at the maximum likelihood estimate for $\theta$, and the E[…] denotes the expectation value of the function in the brackets. We compared the standard error calculated from the empirical Fisher Information to the marginal posterior standard deviation for all transcripts making it past filtering in the Luo et al. dataset (Additional file 1: Fig. S15). While the approximation was better for transcripts with more reads, as expected, the approximation was always an over-estimation of uncertainty for transcripts with low read counts. This property was desirable at it ensured that the MLE implementation was more conservative than either of the other implementations

*Linear modeling of replicate variability trend*

Like the MCMC implementation, the MLE implementation stabilizes uncertainty estimates by using the replicate variability trend in samples from the same experimental condition. First, the total uncertainty is calculated for each transcript. This means combining the estimated replicate variability for each transcript with the per-replicate $\mathrm{logit}(\theta)$ uncertainties. Rather than continuing to work with $\mathrm{logit}(\theta)$ estimates, bakR converts the the $\mathrm{logit}(\theta)$ estimates and uncertainties to log($k_{\deg}$) estimates and uncertainties. The delta approximation is used to convert the $\mathrm{logit}(\theta)$ estimate to the log($k_{\deg}$) estimate, and the uncertainty of the log($k_{\deg}$) estimate is calculated using the Fisher Information of the model with a log($k_{\deg}$) parameterization. Assuming that each estimate for log($k_{\deg}$) is drawn from a normal distribution with standard deviation equal to the estimated replicate variability, then the total standard deviation of the average log($k_{deg}$) estimate is:

$$\sigma_{\mathrm{tot}} =\sqrt{\sum_{i = 1}^{\mathrm{nreps}} \hat{\sigma^{2}} + {\sigma_{\mathrm{MLE},i}}^{2}}$$

where $\hat{\sigma^{2}}$ is the estimated replicate variability (i.e., standard deviation of the replicate log($k_{\deg}$) estimates, and ${\sigma_{MLE,i}}^{2}$ per-replicate log($k_{\deg}$) uncertainty in the i-th replicate. Next, transcripts are binned according to their total read counts across all replicates of the same experimental condition. Third, the read counts and total uncertainties of transcripts in each bin are averaged. Finally, a linear regression of log_10_(read counts) vs. ln($\sigma_{\mathrm{tot}}$) is performed to obtain slopes and intercepts for the replicate variability trend in each experimental condition. The regression fit is used in the next step to regularize the final uncertainty estimates. (4)

*Approximate partial pooling*

The hierarchical model described in the MCMC implementation is approximated using the analytical solution to a simple Bayesian model: the Normal distribution model with known standard deviation and unknown mean. With a conjugate prior (normal distribution with mean *μ*_prior_ and precision *ξ*_prior_; precision is the inverse of the variance. Parameterizing the normal distribution according to the precision rather than the variance or standard deviation makes the analytical posteriors more mathematically succinct), the posterior mean is:

$$\mu_{\mathrm{post}} = \bar{\mu}\frac{n*\xi_{o}}{n*\xi_{o} + \xi_{\mathrm{prior}}} + \mu_{\mathrm{prior}}\frac{\xi_{\mathrm{prior}}}{n*\xi_{o} + \xi_{\mathrm{prior}}}$$

where $\xi_{o}$ is the known precision of the normal distribution used to model replicate variability. This model is used is twice, first to regularize the final uncertainty estimate and second to regularize the average log($k_{\deg}$) estimates. For uncertainty regularization, $\bar{\mu}$ is $\sigma_{\mathrm{tot}}$ for the transcript, $\mu_{\mathrm{prior}}$ is the regression model’s prediction of the replicate variability given the transcripts raw sequencing depth, $\xi_{o}$ is average variation of all transcript’s replicate variability about the linear trend, n is the number of replicates, and $\xi_{\mathrm{prior}}$ is the linear regression standard error. For regularizing the average log($k_{\deg}$) estimates, $\bar{\mu}$ is the weighted sample average of the log($k_{\deg}$) estimates, weighted by the estimate precisions, $\mu_{prior}$ is the average log($k_{\deg}$) estimate, averaged across all samples from the same experimental condition, $\xi_{o}$ is ${\exp(\mu_{\mathrm{post}})}^{-2}$ with $\mu_{\mathrm{post}}$ from the uncertainty regularization, and $\xi_{\mathrm{prior}}$ is inverse square of the total standard deviation of all log($k_{\deg}$) estimates in a given experimental condition.

The approximate posterior log($k_{\deg}$) estimates and uncertainties are then used to calculate $L2FC(k_{\deg})$estimates and uncertainties. Assuming independence of the log($k_{\deg}$) estimates from two different experimental conditions, the $L2FC(k_{\deg})$ estimate is the difference of the log($k_{\deg}$) estimates (experimental condition minus reference condition) and the uncertainty is the square root of the sum of the squares of the uncertainties of the two log(*k*_deg_) estimates.

**Hybrid implementation details**

The Hybrid implementation combines features of the MLE and MCMC implementations. The replicate logit($\theta$) estimates and standard errors from the MLE implementation are passed as data to a hierarchical Stan model nearly identical to that described in the MCMC implementation section, but without the mixture model. The $\mathrm{logit}(\theta)$ standard errors are used to model the estimated logit$(\theta)$ as being drawn from a normal distribution centered on a “true” average logit($\theta)$ with standard deviation equal to the standard error. Mathematically, the model can thus be described as:

$$\mathrm{logit}(\theta_{s,t}^{\mathrm{err}}) \sim\mathrm{Normal}(\mathrm{logit}(\theta_{s,t}), \mathrm{se}_{s,t})$$

$$\mathrm{logit}(\theta_{s,t})\sim\mathrm{Normal}\left( \theta_{e,t},\sigma_{e,t}^{\mathrm{rep}} \right)$$

$$\theta_{e,t} \sim\mathrm{Normal}(\theta_{e},\sigma_{e})$$

$$\ln(\sigma_{e,t}^{\mathrm{rep}}) \sim\mathrm{Normal}(\mu_{e}^{\mathrm{trend}},\sigma_{e}^{\mathrm{trend}})$$

$$\mu_{e}^{\mathrm{trend}} \sim a_{e}*\log_{10}({\bar{\mathrm{reads}}}_{e,t}) +b_{e}$$

where $\mathrm{logit}(\theta_{s,t}^{\mathrm{err}})$ is the logit($\theta)$ estimate provided by the MLE implementation, and $\mathrm{se}_{s,t}$ is the estimate’s standard error (estimated using the Fisher Information as described in **MLE implementation details**). The priors and parameterizations used are identical to that described in **Methods** for the MCMC implementation.

**Other model implementations and naming rationale**

The other models considered for use in bakR were pulseR’s complete pooling negative binomial (CPNB) model and Schofield et al./GRAND-SLAM’s non-hierarchical Poisson mixture model (NHPMM). We refer to pulseR’s model as a complete pooling negative binomial model because it assumes that the negative binomial dispersion parameter is equal for all transcripts (i.e., it completely pools the dispersion parameter estimates). We refer to Schofield et al./GRAND-SLAM’s mixture models as non-hierarchical because no partial pooling of *θ* estimates is performed by this model.

For the CPNB model, we used pulseR’s implementation for both parameter and confidence interval estimation. The boundaries for parameter estimation were set as follows: dispersion parameter (size) between 1 and 10^6^_,_ ln(mean) (mu) between -2.3 and 13.82, log(degradation rate constant) (d) between -7.6 and 2.3, and normFactors set to 1.

For the NHPMM model, we used Stan as with Schofield et al.’s original implementation and the MCMC implementation of the novel hierarchical model. The generative model is identical to the novel hierarchical Poisson mixture model, minus the hyperparameters that make the novel model hierarchical. Both the NHPMM and CPNB implementations rely on bakR for preprocessing the analyzed data (i.e., filtering out difficult to analyze transcripts and getting data into a form that can be passed to the statistical models). The CPNB implementation also required its own additional preprocessing function to make the processed data pulseR-compatible.

**Statistical inference pragmatism**

Much has been written regarding the fraught interpretation of p values and their overuse in the sciences (5). While some of these problems are partially remedied in high-throughput datasets (e.g., Benjamini-Hochberg multiple test adjusted p values are more closely interpretable as posterior probabilities that a transcript is null given the data rather than the more convoluted inverse probability (6)), complaints such as the unrealistic nature of exactly 0 null hypotheses remain. We believe that despite these limitations, null hypothesis testing is a pragmatic strategy by which to rigorously order a list of transcripts and select the transcripts with the most convincing kinetic differences, or to conclude that no such high confidence differences exist. That being said, one additional feature implemented in bakR is the ability to use more realistic compound null hypotheses. More specifically, users can test the null hypothesis that the magnitude of the true $L2FC(k_{\deg})$ is less than some threshold (7). Testing against a magnitude threshold better aligns with the idea of testing against the hypothesis of no biological significance. Thus, while it is still difficult to objectively choose a biological significance threshold, the flexibility of bakR allows it to address some of the concerns of simple null hypothesis statistical testing.

**References:**

1.	Gelman A. Bayesian data analysis. Third edition. ed. Boca Raton: CRC Press; 2014. xiv, 661 pages p.

2.	Team RC. R: A language and environment for statistical computing. R Foundation for Statistical Computing, Vienna, Austria. 2022;<https://www.R-project.org/>.

3.	Rice JA. Mathematical statistics and data analysis. Monterey, Calif.: Brooks/Cole Pub. Co.; 1988. xx, 594 p. p.

4.	Benjamini Y, Hochberg Y. Controlling the False Discovery Rate - a Practical and Powerful Approach to Multiple Testing. J R Stat Soc B. 1995;57(1):289-300.

5.	Wagenmakers EJ. A practical solution to the pervasive problems of p values. Psychon B Rev. 2007;14(5):779-804.

6.	Storey JD. The positive false discovery rate: A Bayesian interpretation and the q-value. Ann Stat. 2003;31(6):2013-35.

7.	McCarthy DJ, Smyth GK. Testing significance relative to a fold-change threshold is a TREAT. Bioinformatics. 2009;25(6):765-71.
